# Vocal ontogeny in *Mus musculus*

**DOI:** 10.1101/2025.03.24.644972

**Authors:** Nicole M. Pranic, Rhea Singh, Joclin Rabinovich, Delia Ferry, Katherine A. Tschida

**Author notes:** Corresponding Author: Katherine A. Tschida.

## Abstract

Infants of many species produce distress calls when in need of parental care. As they mature and gain independence from caregivers, juveniles stop producing infant calls and begin producing adult-like vocalizations in a variety of species-typical contexts. Neonatal mice (*Mus musculus*) produce “isolation” ultrasonic vocalizations (USVs) when separated from the nest, and in contrast, adolescent and adult mice produce “social” USVs during interactions with conspecifics. Although this transition in vocal behavior is known to occur, its developmental timing remains poorly described. In the current study, we performed longitudinal measurements of mouse USVs to establish the age at which juvenile mice begin producing social USVs. Every 4 days (postnatal day (P)12-P28), USVs were first recorded from individual mice in solo sessions to establish rates of isolation/spontaneous USVs and then were recorded again during interactions between same-sex and opposite-sex pairs of age-matched, socially novel (non-littermate) C57BL/6J mice. We report that juvenile mice begin producing social USVs earlier than previously reported, at ages P20-P24. Moreover, although these early social USVs are produced at low rates compared to adult social USVs, they are nonetheless temporally coordinated with active social interaction. Our characterization of the timing of the transition from isolation USVs to social USVs forms a foundation to investigate brain mechanisms that enable the context-dependent regulation of vocal communication over development.

## Introduction

Vocal communication is an important component of social behavior in humans and other animals. Although many animals produce vocalizations across the lifespan, the cues and contexts that regulate vocal communication can differ dramatically during different developmental stages. Infants of many species vocalize when hungry, cold, or isolated from caregivers (Awam et al., 2011; Hahn & Schanz, 2002; Lester & Boukydis, 1985; Monticelli et al., 2004; Newman, 2007; Weary et al., 1997; Zippelius & Schleidt, 1956). These infant distress vocalizations are crucial to the survival of young altricial animals because they attract caregiver attention (Haskins, 1977; Noirot, 1972; Smith, 1976; Soltis, 2004; Symmes & Biben, 1985). As juveniles mature and become less dependent on caregivers, they stop producing infant distress vocalizations and begin producing adult-like vocalizations in a variety of contexts, including during interactions with social partners and in response to social cues (Campbell et al., 2014; Grimsley et al., 2011; Oller et al., 2013; Prat et al., 2015). Characterizing the developmental timing of this transition in vocal behavior is a key step to understanding the brain mechanisms that enable the context-dependent regulation of vocal communication over development.

Laboratory mice (*Mus musculus*) are a valuable model organism to study developmental transitions in vocal communication. Adult female and male mice produce ultrasonic vocalizations (i.e., USVs) when interacting with same-sex and opposite-sex conspecifics (i.e., social USVs) (Gourbal et al., 2004; Maggio & Whitney, 1985; Moles et al., 2007; Neunuebel et al., 2015; Nyby 1983; Portfors & Perkel, 2014; Warren et al., 2018; White et al., 1998; Zhao et al., 2021, 2024). Neonatal mouse pups, on the other hand, produce USVs in response to isolation from the nest and/or decreases in body temperature (isolation USVs) (Zippelius & Schleidt, 1956). Isolation USVs promote the survival of mouse pups, as they elicit a maternal retrieval response (Ehret & Haack, 1984; Noirot, 1972). As pups gain the ability to locomote and thermoregulate independently, rates of isolation USVs decline and reach near-zero at approximately postnatal day (P) 20 (Castellucci et al., 2018; Grimsley et al., 2011; Pranic et al., 2022). Previous work has shown that by early adolescence (∼P30), both opposite and same-sex pairs of age-matched C57BL/6J mice produce USVs during social interactions (Panksepp et al., 2007; Peleh et al., 2019; Scattoni et al., 2013), rates of which are correlated with the amount of time pairs spend socially investigating each other (Panksepp et al., 2007). Whether younger mice produce social USVs during interactions with conspecifics remains untested. Furthermore, whether the earliest social USVs are coordinated with social investigation, as is the case in P30 adolescents (Panksepp et al., 2007) and in adults (Chabout et al., 2012; Heckman et al., 2017; Moles et al., 2007; Zhao et al., 2021) remains unknown.

To address these questions in the current study, we performed longitudinal measurements of mouse USVs to establish the age at which juvenile mice begin producing social USVs. At each age (every 4 days, P12-P28), USVs were first recorded from individual mice in solo sessions to establish rates of isolation and/or spontaneous USVs and then were recorded again after pairs of age-matched, socially novel (non-littermate) C57BL/6J mice were placed together and allowed to interact freely. To test whether early social USVs were coordinated with social investigation or other social behaviors, we also measured non-vocal social behaviors from video recordings of interaction sessions.

## Materials and Methods

Further information and requests for resources should be directed to the corresponding author, Katherine Tschida.

### Ethics Statement

All experiments and procedures were conducted according to protocols approved by the Cornell University Institutional Animal Care and Use Committee (protocol #2020-001).

### Subjects

Female (n = 28) and male (n = 30) C57BL/6J mice (Jackson Laboratories, 000664) were kept on a 12h:12h reversed light/dark cycle and given *ad libitum* chow and water for the duration of the experiment. Mice were housed with their siblings and their mother until weaning at P21, with the exception of one group (n = 3) of females that were housed with their siblings and both of their parents until weaning at P21. Following weaning, mice were housed with their same-sex siblings. To enable individual identification, mice were ear-notched at P11.

Data from adult C57BL/6J mice (> 8 weeks of age) were collected as part of a separate study (Zhao et al., 2021). Those mice were weaned and housed as described above but were not ear-notched until adulthood.

### Study design

To characterize the developmental transition from isolation USVs to social USVs, we longitudinally recorded USVs and non-vocal social behaviors of age-matched, non-sibling pairs of mice (i.e., mice from different litters born 1 day or less apart) during 10-minute-long interaction sessions at P12, P16, P20, P24, and P28. We longitudinally recorded behaviors of pairs of male-female (MF), female-female (FF), and male-male (MM) mice, and any given mouse was only ever used in a single social context (MF, FF, or MM). Due to experimenter error, one cohort of FF pairs of mice (n = 3) was tested at P23 rather than P24. Mice were paired with non-siblings during interaction sessions but were not paired with the same non-sibling social partner in consecutive interaction sessions. Prior to each interaction session, we recorded USVs from each individual mouse alone for 10 minutes in a cage with clean bedding (solo sessions).

During interaction sessions, pairs of mice were placed in a lidless acrylic cage with clean bedding that was in turn placed inside a high-walled, lidless acrylic chamber inside a sound-attenuating recording chamber (Med Associates). The recording chamber was equipped with an ultrasonic microphone (Avisoft), an infrared light source (Tendelux), and a webcam (Logitech, with the infrared filter removed to enable video recording under infrared lighting). Prior to the start of the experiment, subject mice had only ever interacted with their siblings and parents. The sample sizes within each social context are as follows: n = 10 MF pairs; n = 9 FF pairs; n = 10 MM pairs of mice. An interaction session of one pair of MM P28 mice was excluded from analysis due to one of the mice jumping out of the cage during the trial.

Data from 30-minute-long interaction sessions between pairs of group-housed non-age-matched adult mice included in this manuscript were collected as part of a previously published study (Zhao et al., 2021). On the day of the behavior measurements, the subject mouse was transferred in its home cage to a sound-attenuating recording chamber equipped for video and audio recording as described above. Siblings of the subject mouse were removed from the home cage and transferred to a clean cage for the duration of the interaction session. A novel, unfamiliar group-housed visitor mouse (male or female depending on the social context) was then placed in the resident mouse’s home cage for 30 minutes, and video and audio recordings were made. Visitors were marked with a small spot of acrylic paint to facilitate identification in videos. Measurements of vocal and non-vocal behaviors were made in three social contexts: female residents with female visitors (n = 31 FF trials), male residents with male visitors (n = 20 MM trials), and male residents with female visitors (n = 22 MF trials).

### USV recording and detection

USVs were recorded using an ultrasonic microphone (Avisoft, CMPA/CM16), connected to an Avisoft recording system (UltrasoundGate 116H, 250 kHz sample rate). Because USV rates were low, particularly in P20-P28 mice, and to ensure the highest accuracy of USV detection, we manually annotated all USVs from spectrograms generated from audio files using custom MATLAB codes. USVs recorded during interaction sessions between adult mice were also manually annotated (Zhao et al., 2021). USV rates reported in interaction sessions represent the total number of USVs produced by each pair.

### Non-vocal behavior analysis

Trained observers scored the following behaviors from webcam recordings of interaction sessions: (1) not interacting, (2) active social interaction (sniffing, following, grooming of partner, or mounting), (3) huddling (pairs of mice in side-by-side physical contact for more than 1 second without engaging in active social interaction as defined above), and (4) fighting (attacking and/or biting). We did not observe any instances of mounting or fighting among P12-28 mice. We did not observe any instances of huddling among adult mice. Non-vocal behaviors were scored as mutually exclusive continuous events. A bout of active social interaction was defined as a period of active social interaction that was sustained for at least 1s, and distinct bouts of active social interaction were separated by a period of at least 1s in which mice stopped interacting.

### Statistical analyses

Details of the statistical analyses used in this study are included in Table S1. To determine whether to use parametric or non-parametric statistical models and tests for a given comparison, we examined the normality of the residuals for the relevant data distributions (determined by visual inspection of plots of z-scored residuals; cases in which residuals diverged notably from the 45-degree line of a normal distribution were deemed non-normally distributed). All p-values for pairwise comparisons were corrected for multiple comparisons using the Tukey honestly significant difference (HSD) test. No statistical methods were used to pre-determine sample size. All statistical analyses were carried out using R 4.3.0 (R Core Team, 2023) and R Studio 2023.03.1+446 (Posit team, 2023).

### Data availability

All source data generated in this study will be deposited in a digital data repository, and this section will be modified prior to publication to include the persistent DOI for the dataset.

## Results

### Mice begin producing social USVs by postnatal day 20-24

To establish the earliest age at which we could reliably detect the production of social USVs by juvenile mice, we compared rates of USVs produced by juveniles that were recorded alone (solo sessions) to rates of USVs produced by the same juveniles that were subsequently given a social interaction with an age-matched, socially novel (non-littermate) juvenile (interaction sessions; Fig. 1A). These solo and interaction sessions were conducted longitudinally every 4 days from P12-P28, and a given mouse only ever participated in interaction sessions in a single social context (male-female interactions (MF), female-female interactions (FF), or male-male interactions (MM); n = 9-10 pairs for each age and social context). Because some USVs produced by juvenile mice were low amplitude, we manually annotated USVs from spectrograms of audio files to obtain USV counts from solo and interaction sessions.

**Figure 1.**
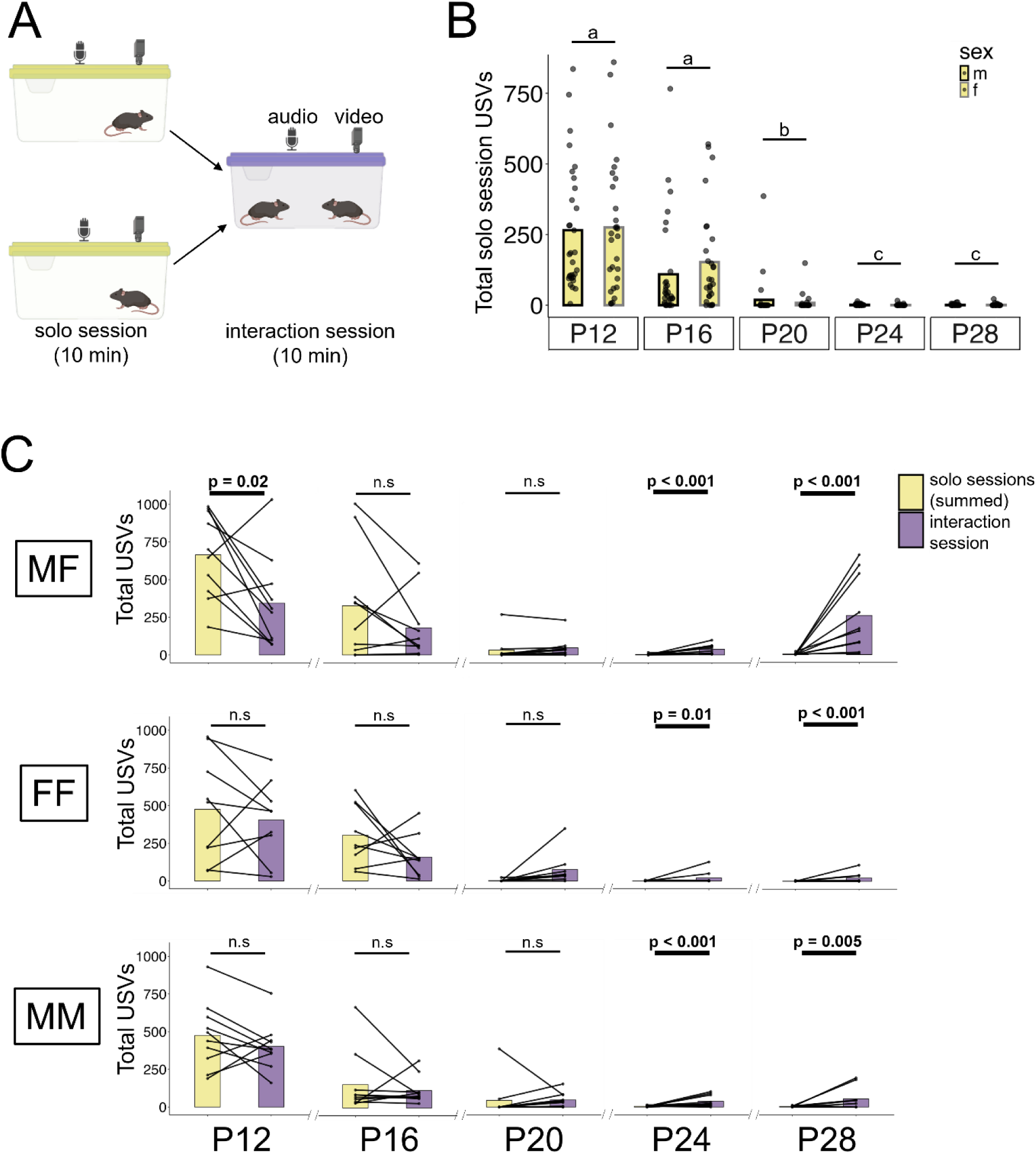
Mice produce social USVs by postnatal day 24 during both same-sex and opposite-sex interactions. (A) Schematic shows recording of USVs in solo sessions, followed by recording of USVs and non-vocal behaviors in interaction sessions. (B) Counts of solo session USVs are shown for males (black outlines) and females (gray outlines), aged P12-P28. Different letters indicate statistically significant differences. (C) Total summed solo session USVs (yellow bars) vs. total interaction session USVs (purple bars) are shown for pairs of age-matched mice, aged P12-P28. MF = male-female, FF = female-female, MM = male-male. The experimental design schematic included in (A) was created using icons generated with BioRender (https://app.biorender.com/illustrations/67de0ce2267e5f30068a1873?slideId=4d8be0e1-a899-424d-99e2-9d68284f8470).

Consistent with prior findings that rates of mouse isolation USVs decline dramatically by the third postnatal week (Castellucci et al., 2018; Grimsley et al., 2011; Pranic et al., 2022), we found that solo session USV rates declined from P12 to P28 in both female and male mice, reaching very low rates by P20 and reaching near-zero rates by P24 (Fig. 1B; negative binomial generalized linear mixed model (glmm-nb), p < 0.01 for P12/P16 vs. P20, P12/16 vs. P24/28, and for P20 vs. P24/28; see Table S1 for full details of statistical analyses).

We reasoned that when juvenile mice begin producing social USVs, one would expect the total USVs produced during interaction sessions to exceed those produced during solo sessions. For each age and social context, we therefore compared the total USVs produced during each interaction session to the sum of the total USVs produced by each of the two mice in their respective solo sessions. For mice aged P12-P20, we found that total interaction session USVs were either equal to or less than the summed total of USVs produced during solo sessions, and this was true across all three social contexts (Fig. 1C). In contrast, starting at P24, we found that mice produced significantly more USVs during interaction sessions than during solo sessions (Fig. 1C; glmm-nb, p < 0.05 for all P24 and P28 summed solo vs. interaction comparisons). Although the difference between summed solo and interaction session USV rates is significant for both P24 and P28 pairs, we note that rates of USVs produced by P24 and P28 pairs tend to be lower than those produced by group-housed adults engaged in same-sex and opposite-sex interactions (Fig. S1). These analyses support the conclusion that by P24, juvenile mice begin producing social USVs during interactions with both same-sex and opposite-sex juvenile social partners.

When considering isolation USV rates from P20 juveniles, we noticed that although most P20 juveniles produced near-zero USVs in their solo sessions, there were a handful of P20 juveniles that continued to produce modest rates of isolation USVs (Fig. 1B; n = 3 P20 females and n = 3 P20 males produced > 20 USVs in solo sessions). In other words, there may be individual differences in developmental trajectories, whereby most P20 juveniles have ceased producing isolation USVs, while a small number continue doing so. We wondered whether the inclusion of interaction trials that included these juveniles might have obscured our ability to detect social USV production from the other P20 pairs in our dataset. To test this idea, we re-considered the P20 dataset after excluding the 5 pairs (n = 2 MF, n = 1 FF, and n = 2 MM) that included individual P20 mice that produced >20 USVs in their solo sessions. Analysis of this modified dataset revealed that P20 MF and P20 MM pairs produced significantly more USVs during interaction sessions than during solo sessions, with a similar although non-significant trend observed in P20 FF pairs (Fig. 2; lmm, p < 0.05 for MF and MM summed solo vs. interaction comparisons, p = 0.056 for FF summed solo vs. interaction). In summary, our findings support the conclusion that juvenile mice begin producing social USVs at younger ages than previously reported (P20-24), in a manner that temporally coincides with a dramatic decline in isolation USV production.

**Figure 2.**
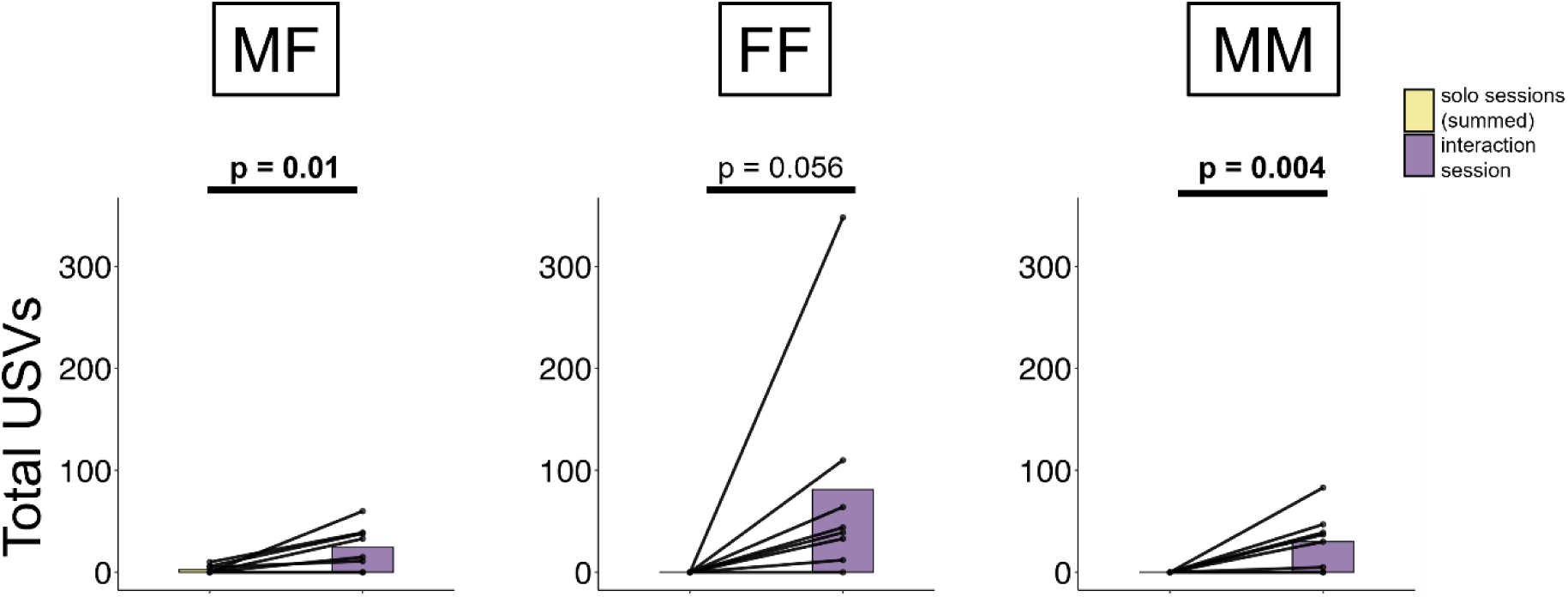
Evidence that some mice produce social USVs by postnatal day 20. (A) Total summed solo session USVs (yellow bars) vs. total interaction session USVs (purple bars) are shown for pairs of age-matched mice, aged P20. These analyses were conducted after first excluding data from n = 5 pairs that included individual P20 mice that produced > 20 USVs in their solo sessions. MF = male-female, FF = female-female, MM = male-male.

### Description of non-vocal social behaviors produced by juvenile mice

As a first step toward testing whether early social USVs are coordinated with non-vocal social behaviors, we characterized the non-vocal social behaviors produced by juveniles at different ages and quantified developmental changes in the time spent engaged in these behaviors. We found that pairs of juvenile mice engaged in active social interactions (sniffing, following, or grooming of the other mouse) as well as in huddling (side-by-side physical contact not marked by sniffing, locomotion, or social grooming). Although time spent in active social interaction tended to increase from P12 to P28, this change was not significant for any of the social contexts (Fig. 3A, blue bars; lmm, p > 0.05 for MF, FF, and MM). Even though total time spent in active social interaction did not change significantly over the ages we considered, we found that the temporal organization of social interaction bouts changed with age. Specifically, we found that the mean duration of social interaction bouts decreased with age, while the mean total number of social interaction bouts increased with age (Fig. 3B-C; different letters indicate statistically significant differences; glmm-nb to analyze 3B, lmm to analyze 3C). Time spent huddling peaked at P16, after which it declined and reached low rates by P24 (Fig. 3A, pink bars). Finally, time spent not interacting reached its lowest value at P16 (at the same age at which huddling peaked) but occurred at similar rates across the other measured ages (Fig. 3A, gray bars). Neither mounting nor fighting were observed in juvenile interaction sessions.

**Figure 3.**
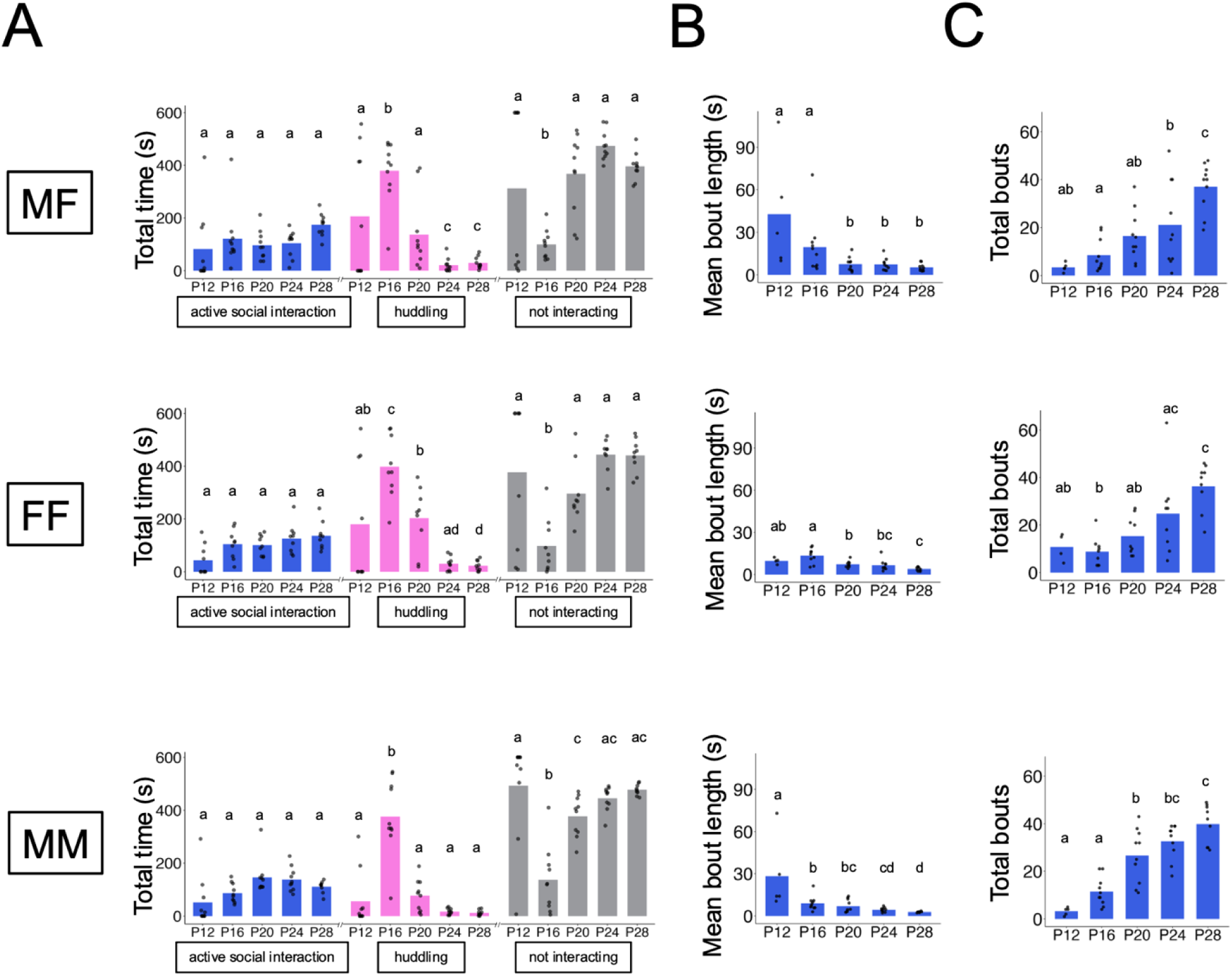
Description of time spent in non-vocal social behaviors. (A) The total time (out of 600 total seconds) spent engaged in different non-vocal behaviors is shown for age-matched pairs, plotted by age and by social context. Blue = active social interaction (sniffing, following, or grooming of the other mouse), pink = huddling, gray = not interacting. Different letters indicate statistically significant differences. (B) Mean length of active social interaction bouts is plotted by age and by social context. (C) Same as (B), for total bouts of active social interaction. MF = male-female, FF = female-female, MM = male-male.

### Early social USVs are coordinated with non-vocal social behaviors

In group-housed adult mice, USV production is positively correlated with total social interaction time only in certain social contexts; namely, during male-female and female-female interactions, but not during male-male interactions (Chabout et al., 2012; de Chaumont et al., 2021; Zhao et al., 2021). A previous study measuring social interactions between age-matched P30 mice reported that the total number of USVs was positively correlated with social interaction time (defined as the sum of time spent sniffing, following, or social grooming, as in the current study), although this relationship was not assessed separately for trials from opposite-sex vs. same-sex pairs (Panksepp et al., 2007). To test whether social USV production and social interaction time are positively related across the three juvenile social interaction contexts, we compared total USVs to the total time spent in active social interaction for interaction trials from MF, FF, and MM trials (Fig. 4). Given that P24 is the youngest age in our dataset at which social USV production was statistically significant across all social contexts (Fig. 1C), we focused these and subsequent analyses on P24 and P28 pairs. In P24 pairs, total USVs and total active social interaction time were uncorrelated for all social contexts (Fig. 4; linear regression, p > 0.05 for all). Notably, a positive and statistically significant relationship emerged at P28 for both opposite-sex and same-sex pairs of mice (Fig. 4; p < 0.05 for all). For ease of comparison, our previous behavioral data collected from group-housed adult interaction trials are re-plotted here, showing that USV production is significantly correlated with total interaction time only in MF and FF pairs, but not in MM pairs (Fig. 4; data replotted from Zhao et al., 2021). In summary, a positive relationship between total USVs and social interaction time is not present at P24, emerges at P28 for all social contexts, and then persists into adulthood for only male-female and female-female interactions.

**Figure 4.**
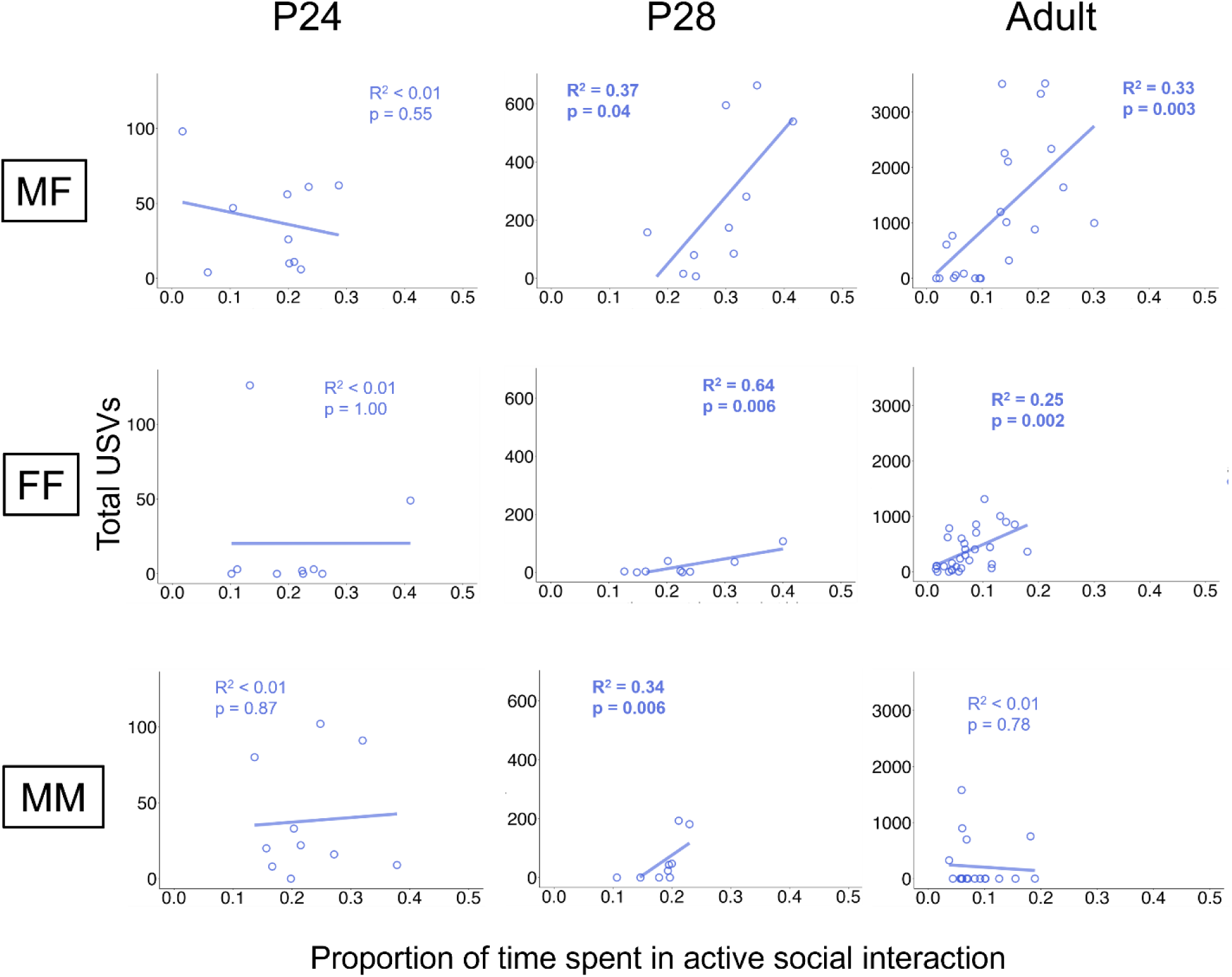
Relationship of total USVs to time spent in active social interaction. Scatterplots show the proportion of total time (out of 600 total seconds) spent in active social interaction vs. total USVs for interaction sessions for P24, P28, and adult mice during same-sex and opposite-sex interactions. MF = male-female, FF = female-female, MM = male-male.

Group-housed adult male-female and female-female pairs of mice produce the majority of their USVs as they engage in active social interaction (de Chaumont et al., 2021; Zhao et al., 2021), and the positive correlation that exists between total USVs and total active social interaction time for P28 pairs suggests that USV production and social interaction are also temporally coordinated at this earlier age. Although total USVs are not well related to total interaction time in P24 pairs, it is important to note that social USV rates at this age are very low (Fig. 1C; see also Fig. S1 for a comparison of USV rates during interactions for P24 vs. P28 vs. adult pairs). Thus, one possibility is that many P24 social USVs are produced during periods of active social interaction, but not all periods of social interaction are accompanied by USV production. More generally, we wanted to understand what non-vocal behaviors were being produced by P24 and P28 mice as they produced social USVs. To answer these questions, we assigned a non-vocal behavior code to each USV produced during P24 and P28 interaction trials, which corresponds to the non-vocal behavior that the pair was performing when that USV was produced (active social interaction, huddling, or not interacting). Because there is substantial variability within and across ages in the proportion of time that pairs spent engaged in different non-vocal behaviors (Fig. 3A), we divided the number of USVs that each pair produced during different non-vocal behaviors by the total time that the pair spent engaged in each non-vocal behavior (USVs per second of behavior; Fig. 5). This analysis of normalized USV rates revealed that P24 and P28 mice tend to produce most of their USVs while engaged in active social interaction (Fig. 5). This trend was statistically significant for P28 MF trials and P28 MM trials, and we observed similar, though non-significant trends for P24 trials and for P28 FF trials (lmm, p < 0.001 for all P28 MF comparisons, p < 0.01 for all P28 MM comparisons, p > 0.05 for all other comparisons).

**Figure 5.**
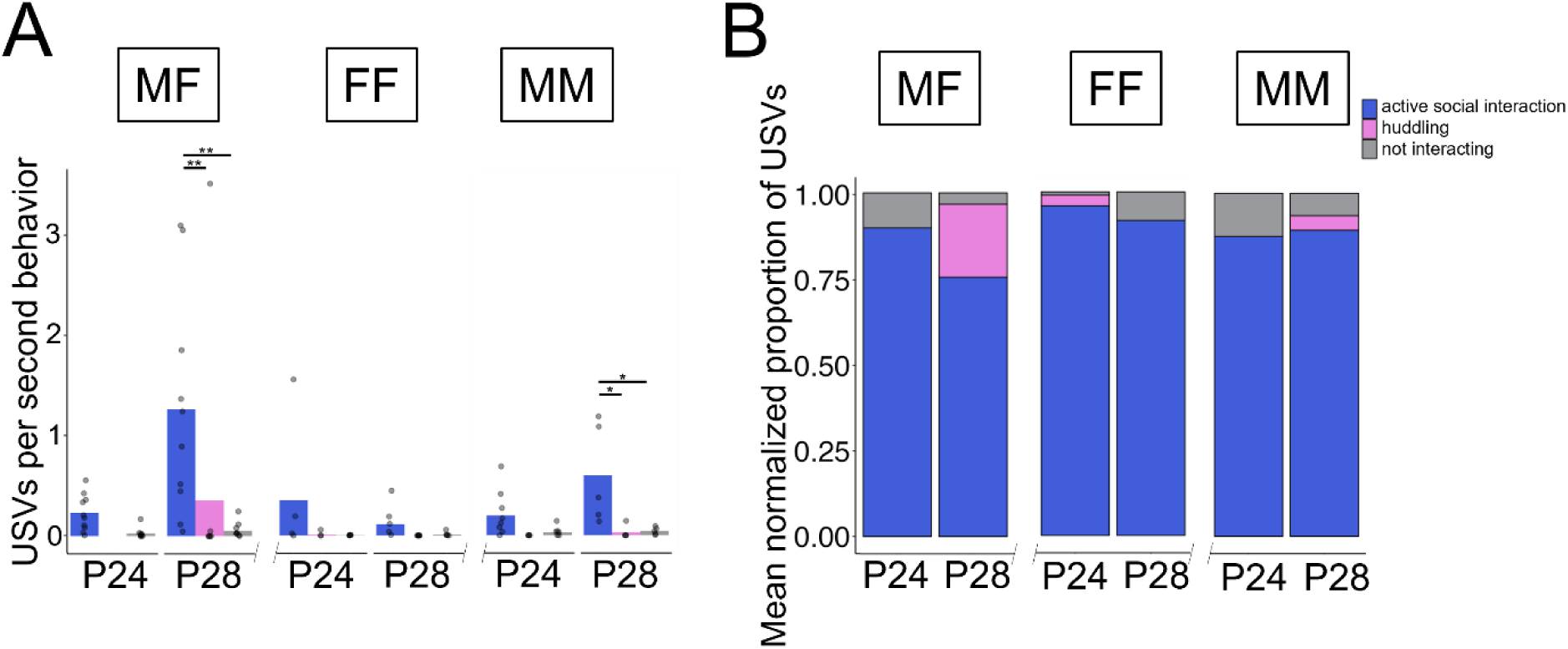
P24 and P28 pairs tends to produce social USVs during active social interaction. (A) The mean number of USVs produced per second of non-vocal behavior (active social interaction, huddling, or not interacting) during interaction sessions of same-sex and opposite-sex P24 and P28 pairs. (B) Stacked bar plot shows the mean proportion of total USVs produced during each non-vocal behavior, normalized by the amount of time each pair spent performing that behavior. MF = male-female, FF = female-female, MM = male-male.

To visually compare the timing of social USV production to the timing of different non-vocal behaviors, we generated ethograms of USV rates aligned with categories of non-vocal behavior produced during representative P24 and P28 trials (Fig. 6A). To facilitate comparisons with adult behavioral data, we also included representative ethograms from adult opposite-sex and same-sex interactions (data replotted from Zhao et al., 2021). These visualizations support the idea that although USV production is often temporally coordinated with active social interaction, not all social interactions are accompanied by USV production, particularly in juvenile interaction trials. Within these ethograms, we also noticed that a substantial proportion of total USV production and total active social interaction seem to take place early on during interaction trials (Fig. 6A). To quantify this observation, we calculated the proportion of total USVs and the proportion of total active social interaction that occurred during 100-second-long bins within P24, P28, and adult interaction trials (Fig. 6B; adult trials were 30 minutes in duration, and here we only considered behavior from the first 10 minutes of those trials). We found that for almost all ages and social contexts, the majority of USVs and the majority of active social interaction occurred during the first 100 seconds of interaction trials, with the exception of MF pairs of adult mice (Figs. 6B & S2). In contrast to the other groups, MF adult pairs exhibited USV production and social interaction that was on average more evenly sustained across the entire 10 minutes (Figs. 6B & S2). Furthermore, although both USV production and active social interaction peaked early on during interaction trials for most ages and social contexts, this effect was in general more pronounced for USV production than for interaction time (Figs. 6B & S2). In summary, we found that early social USVs are temporally coordinated with active social interaction in opposite-sex and same-sex pairs of P24 and P28 mice. We also conclude that both USVs and active social interaction tend to occur at the highest rates early on during interaction trials, and that this temporal bias is particularly pronounced for social USV production in P24 and P28 mice.

**Figure 6.**
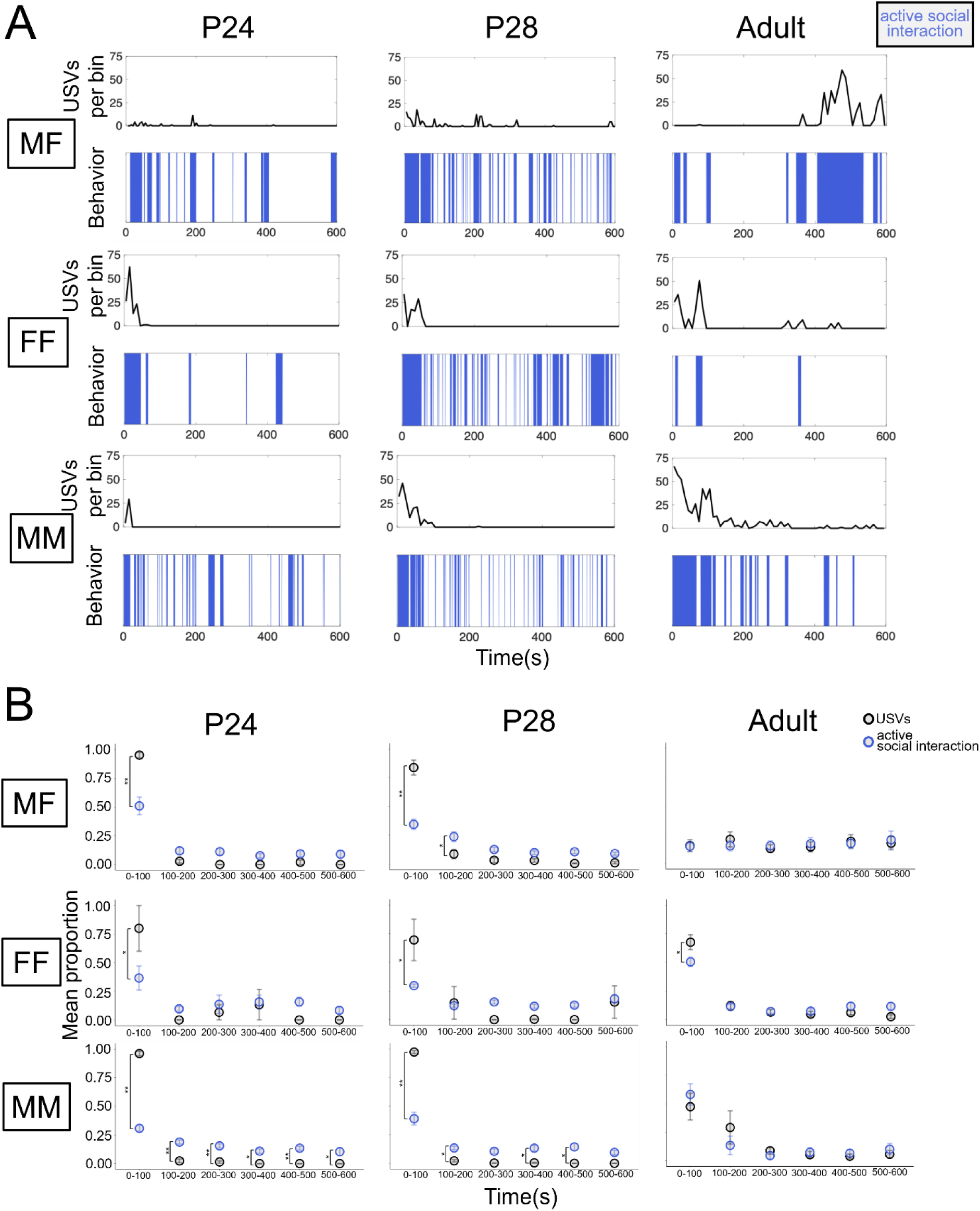
Temporal organization of USVs and active social interaction during interaction sessions. (A) Ethograms are shown for P24, P28, and adult interaction trials. The top half shows USV rate over time (total USVs in each 10s-long bin), and the bottom half shows the occurrence of active social interaction (blue shading) and not interacting (white). Huddling, mounting, and fighting did not occur in the representative trials plotted here. (B) Plots show the proportion of total USVs (black symbols) and the proportion of total active social interaction (blue symbols) occurring in 100s-long bins. Statistical comparisons are shown for the proportion of total USVs vs. the proportion of total active social interaction within each 100s-long bin. MF = male-female, FF = female-female, MM = male-male.

## Discussion

In this study, we describe the ontogeny of mouse social USVs and characterize the coordination of early social USVs with non-vocal social behaviors. We show that mice produce social USVs as early as P20-P24, during interactions with both same-sex and opposite-sex juvenile social partners (Figs. 1 & 2), and that rates of social USVs tend to increase with age (Figs. 1C & S1). With regard to non-vocal social behaviors, we found that although time spent in active social interaction did not increase significantly from P12 to P28, pairs of mice engaged in shorter but more frequent social interaction bouts as they aged (Fig. 3B-C). We found that a positive relationship between total social USVs and time spent in social interaction emerges at P28 for all social contexts, then persists into adulthood for only MF and FF contexts (Fig. 4). Moreover, P24 and P28 mice tended to produce most of their USVs while engaged in active social interaction, further supporting the conclusion that early social USVs of juvenile mice are coordinated with non-vocal social behaviors (Fig. 5). Finally, with the exception of MF pairs of adults, pairs of mice produced the highest rates of USVs and spent the most time engaged in active social interaction during the first 100s of social interaction trials (Figs. 6 & S2).

Previous studies have observed social USVs as early as P30 during interaction trials between unfamiliar pairs of age-matched, same-sex (Panksepp et al., 2007; Scattoni et al., 2013) and opposite-sex (Panksepp et al., 2007) mice. Similar to our findings, Panksepp et al. (2007) found that in pairs of unfamiliar age-matched P30 mice, USV rates correlated with the total time that mice spent in social investigation. In recordings from pairs of same-sex age-matched littermates (ages ranged from P24-P29), Peleh et al. (2019) also observed USVs during social interaction trials. We replicate these results and further provide evidence for the production of social USVs at an even earlier age, in P20 pairs of mice.

One intriguing idea is that social USVs might be produced at even earlier ages than detected in our study. In other words, mice could be producing both isolation and social USVs within a given trial during early development. In our study, we did not find significant differences in summed solo session vs. interaction session USV rates in P12 and P16 pairs, and if anything, interaction session USV rates tended to be lower than summed solo session USV rates at these ages. We cannot rule out the possibility that the presence of a social partner might simultaneously suppress the production of isolation USVs while promoting the production of social USVs, a scenario in which there may be no net increase in USVs from solo to interaction sessions. Future work could test this idea by implementing ultrasonic microphone array recordings (e.g., Sterling et al., 2023; Warren et al., 2018) to determine which mouse in the pair is producing individual USVs and exploring the coupling of individual USVs to nonvocal behaviors of mice during interactions with age-matched conspecifics during early development. However, given our findings that social USVs are produced at relatively low rates by P20-P28 mice, we would anticipate that any social USVs produced prior to P20 would also be produced at correspondingly low rates.

Similar to humans, mice experience developmental changes in numerous social behaviors during adolescence. Following weaning, mouse social interactions gradually become more adult-like (Panksepp et al., 2007), as this period sees the onset of sex-biased aggressive (Terranova et al., 1998) and sexual (Jean-Faucher et al., 1978) behaviors towards conspecifics, as well as an overall increase in affiliative social interactions (Terranova et al., 1993) and the onset of social novelty preference (Bian et al., 2022). The trajectory of these behaviors highlights a critical idea: rather than a single time point, puberty involves the gradual development of adult-like social behaviors (Avendaño et al., 2017; Berenbaum et al., 2015; Blaustein et al., 2016; Foilb et al., 2011; Sisk & Foster, 2004). Our results support this framework of puberty as a gradual process, as we observed gradual increases in rates of social USVs (Figs. 1, 2, & S1), as well as changes in the relationship between USVs and non-vocal social behavior (Fig. 4). Notably, we found that juveniles produce social USVs even prior to the emergence of sex-biased patterns of non-vocal social behaviors such as mounting (MF interactions) and aggression (MM interactions). We also found that, except for MF pairs of adult mice, pairs of mice produced the highest rates of USVs and spent the most time engaged in active social interaction during the first 100s of social interaction trials (Figs. 6 & S2). One potential explanation for this result is that adult males exhibit pronounced and sustained sexual motivation during interactions with females, a type of motivation likely not present in the other social contexts and ages that we considered.

Based on our finding that a subset of P20 mice continue to produce modest rates of isolation USVs (Figs. 1B & S1), as well as our finding that rates of social USVs produced by P20-28 pairs are variable (Figs. 1C & 2), we speculate that there may be considerable individual-to-individual variability in the age at which social USVs begin to be produced. The factors that regulate the developmental timing of the transition from isolation USVs to social USVs, as well as the degree to which this transition is malleable vs. relatively fixed, remain unknown. The answers to these questions would be facilitated by the implementation of microphone array recordings or other methods to enable the assignment of each social USV to an individual signaler. Given that we found evidence for social USV production in pre-weaning P20 mice, we can conclude at least that weaning per se is not a critical driver of the transition from isolation USVs to social USVs, although future work might investigate the degree to which weaning age influences the developmental trajectory of vocal communication after the onset of social USV production.

Our behavioral description of the transition from isolation USVs to social USVs sets the stage for future work to identify the neural circuit changes that underlie this developmental transition in vocal communication. Given prior work that identified midbrain neurons important for gating the production of social USVs in adult female and male mice (Tschida et al., 2019; Ziobro et al., 2024; Malone et al., 2025), an exciting possibility is that functional changes in forebrain inputs to the midbrain vocalization circuit might underlie the ability of different contexts to promote USV production during postnatal development vs. adolescence and adulthood. For example, a recent study found that hypothalamic AgRP neurons, which are sensitive to and regulate food intake in adult mice, are responsive to isolation from the nest but not to milk ingestion or milk deprivation during the first two weeks of postnatal life (Zimmer et al., 2019). With regard to developmental changes in vocal communication, one possibility is that forebrain inputs to the midbrain vocalization circuit gain sensitivity to the social cues of novel partners and thereby begin to promote USV production during social interactions.

More broadly, detailed behavioral descriptions of developmental transitions in social behavior, combined with the plethora of transgenic and viral tools available in mice, can be leveraged to identify developmental changes in neural circuits that underlie adolescent behavioral transitions, aiding in our understanding of brain circuits underlying developmental transitions in social behaviors of both human and non-human animals. It is also important to note that many neurodivergent adolescents experience developmental transitions in various social behaviors differently from their neurotypical peers (Coluccia et al., 2017; Cresswell et al., 2019; McKay et al., 2024; Moody et al., 2022). Uncovering brain mechanisms underlying changes in social behaviors during adolescence could help establish novel strategies aimed at improving difficulties neurodivergent adolescents might experience during these transitional periods.

## Acknowledgements

We thank Frank Drake and other CARE staff for their excellent mouse husbandry, and to Stephen Parry from the Cornell Statistical Consulting Unit for statistical consultation. The experimental design schematic included in Figure 1A was created using icons that were generated with BioRender (https://app.biorender.com/illustrations/67de0ce2267e5f30068a1873?slideId=4d8be0e1-a899-424d-99e2-9d68284f8470).

## Data and Resource Availability

The datasets generated during the current study will be made publicly available, and this section will be updated upon acceptance for publication with the persistent DOI for the dataset.

