## Supplemental Information for "Vocal ontogeny in *Mus musculus*"

**Supplementary Information from Pranic et al., 2025**

**Figure S1. Social USV rates tend to increase over development.** Total USVs produced during interaction sessions are compared for P24, P28, and adult mice during same-sex and opposite-sex interactions.

**Figure S2. More details about temporal organization of USVs and active social interaction during interaction sessions.** (A) Plots show the proportion of total USVs occurring in 100s-long bins. Different letters indicate statistically significant differences. (B) Same as (A), for proportion of total active social interaction.

**Table S1. Full details of statistical analyses.**

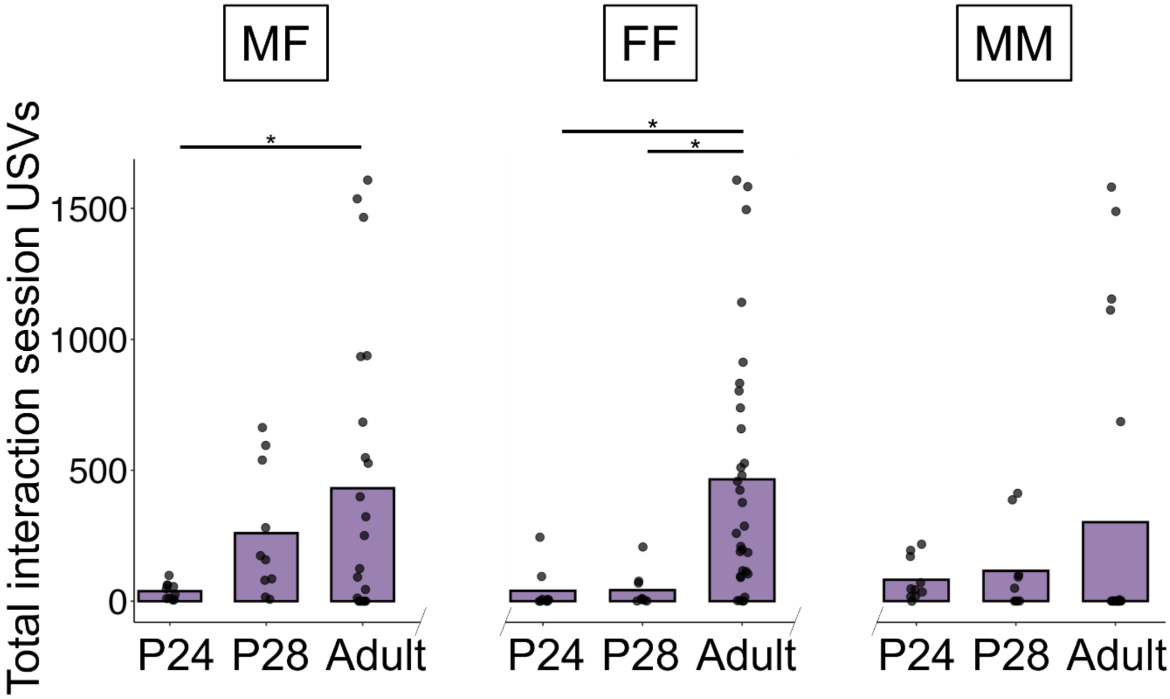

MF
FF
MM
Total interaction session USVs
1500
1000
500
0
*
*
*
P24
P28
Adult
P24
P28
Adult
P24
P28
Adult

**Figure S1**

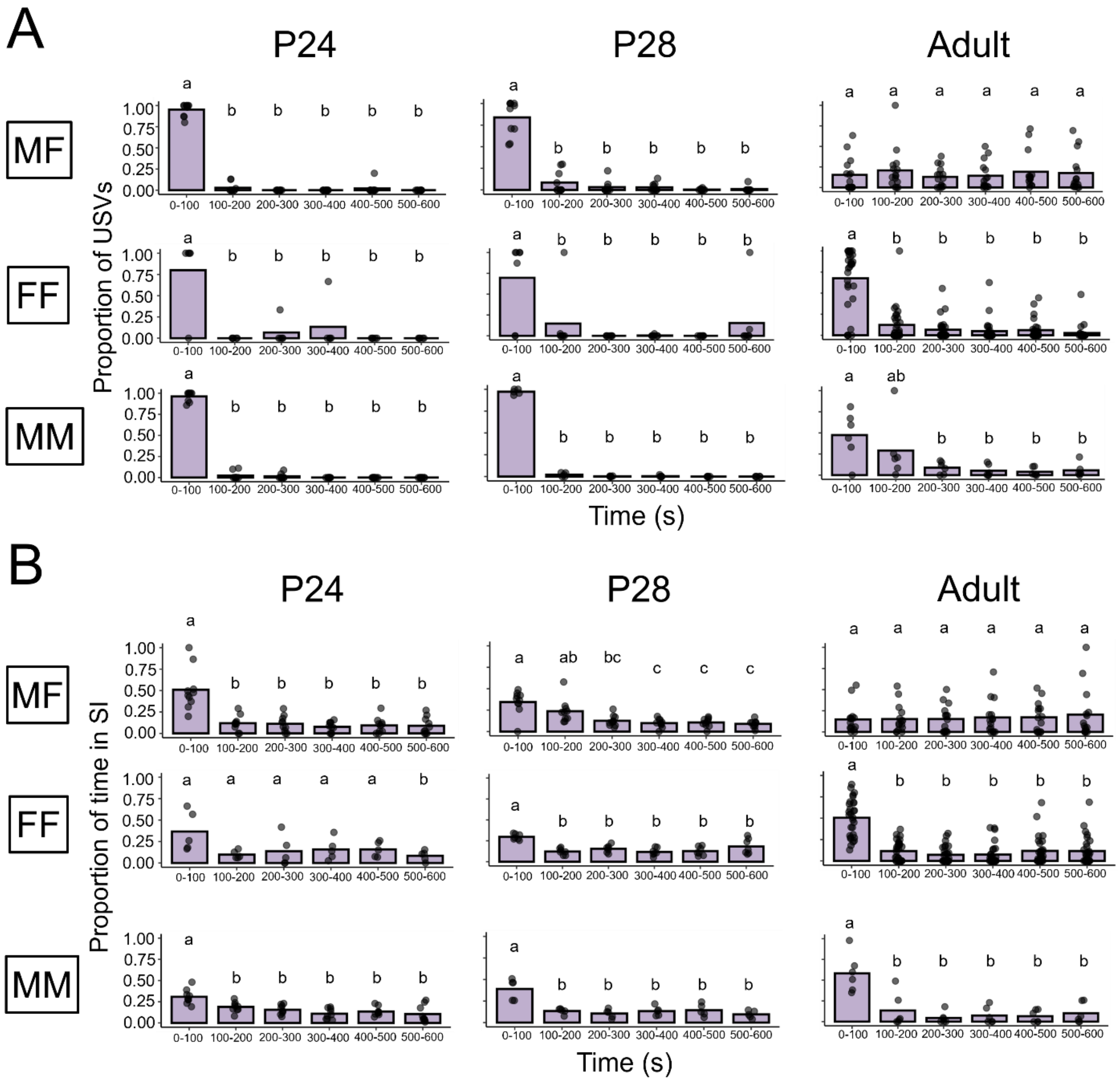
A
P24
P28
Adult
MF
FF
MM
Proportion of USVs
0-100
100-200
200-300
300-400
400-500
500-600
Time (s)
B
P24
P28
Adult
MF
FF
MM
Proportion of time in SI
Time (s)

**Figure S2**

**Table S1**

| Figure | Comparison | Models/tests | Outcome | Descriptive statistics | Notes |
|---|---|---|---|---|---|
| Fig. 1B | solo USV counts x age and sex (cohort and context as random effects) | generalized linear mixed-effects model with negative binomial family (glmm-nb) | $p = 0.21$ (interaction effect between age vs sex)<br>$p < 0.001$ (main effect of age) **<br>$p = 0.93$ (main effect of sex) | Mean ($\bar{x}$) = 265.43, standard deviation (sd) = 219.57 (P12 M)<br>$\bar{x}$ = 275.32, sd = 235.35 (P12 F)<br>$\bar{x}$ = 109.57, sd = 176.18 (P16 M)<br>$\bar{x}$ = 152.57, sd = 174.92 (P16 F)<br>$\bar{x}$ = 18.8, sd = 73.22 (P20 M)<br>$\bar{x}$ = 8.14, sd = 28.94 (P20 F)<br>$\bar{x}$ = 1.8, sd = 3.11 (P24 M)<br>$\bar{x}$ = 0.96, sd = 3.28 (P24 F)<br>$\bar{x}$ = 1.5, sd = 3.11 (P28 M)<br>$\bar{x}$ = 1.5, sd = 4.48 (P28 F) | |
| Fig. 1C (top panel) | MF USV counts x session per age (cohort, mouse identity, dyad identity as random effects) | glmm-nb (P16, P24, P28) linear mixed effects model (lmm; P12, P20) | $p = 0.02$ (P12 main effect of session type)*<br>$p = 0.69$ (P16 main effect of session type)<br>$p = 0.11$(P20 main effect of session type)<br>$p < 0.01$ (P24 main effect of session type)**<br>$p < 0.01$ (P28 main effect of session type)** | $\bar{x}$ = 663.9, sd = 282.23 (P12 solo)<br>$\bar{x}$ = 343.5, sd = 305.45 (P12 interaction)<br>$\bar{x}$ = 326.6, sd = 364.56 (P16 solo)<br>$\bar{x}$ = 180.2, sd = 218.19 (P16 interaction)<br>$\bar{x}$ = 32.9, sd = 83.51 (P20 solo)<br>$\bar{x}$ = 47.4, sd = 67.58 (P20 interaction)<br>$\bar{x}$ = 2.4, sd = 5.08 (P24 solo)<br>$\bar{x}$ = 38.1, sd = 31.53 (P24 interaction)<br>$\bar{x}$ = 3.7, sd = 7.0 (P28 solo) | |

| | | | | |
|---|---|---|---|---|
| | | | | x̅ = 259.8, sd = 248.84 (P28 interaction) |
| Fig. 1C (middle panel) | FF USV counts x session per age (cohort, mouse identity, dyad identity as random effects) | glmm-nb (P16, P24, P28)<br>lmm (P12, P20) | $p = 0.50$ (P12 main effect of session type)<br>$p = 0.08$ (P16 main effect of session type)<br>$p = 0.07$ (P20 main effect of session type)<br>$p = 0.01$ (P24 main effect of session type)*<br>$p < 0.001$ (P28 main effect of session type)** | x̅ = 475.56, sd = 349.25 (P12 solo)<br>x̅ = 404, sd = 257.75 (P12 interaction)<br>x̅ = 304.78, sd = 199.75 (P16 solo)<br>x̅ = 158.89, sd = 142.66 (P16 interaction)<br>x̅ = 2.56, sd = 7.67 (P20 solo)<br>x̅ = 75.78, sd = 106.85 (P20 interaction)<br>x̅ = 0.67, sd = 2 (P24 solo)<br>x̅ = 20.33, sd = 42.69 (P24 interaction)<br>x̅ = 1.11, sd = 2.26 (P28 solo)<br>x̅ = 21.67, sd = 35.51 (P28 interaction) |
| Fig. 1C (bottom panel) | MM USV counts x session per age (cohort, mouse identity, dyad identity as random effects) | glmm-nb (P16, P20, P24, P28)<br>lmm (P12) | $p = 0.26$ (P12 main effect of session type)<br>$p = 0.74$ (P16 main effect of session type)<br>$p = 0.22$ (P20 main effect of session type)<br>$p < 0.001$ (P24 main effect of session type)**<br>$p = 0.005$ (P28 main effect of session type)* | x̅ = 475.30, sd = 220.79 (P12 solo)<br>x̅ = 401.80, sd = 153.80 (P12 interaction)<br>x̅ =155.5, sd = 203.87 (P16 solo)<br>x̅ =115.5, sd = 89.40 (P16 interaction)<br>x̅ = 44, sd = 121.36 (P20 solo)<br>x̅ = 47.5, sd = 47.52 (P20 interaction)<br>x̅ = 5.1, sd = 4,61 (P24 solo)<br>x̅ = 38.1, sd = 37.92 (P24 interaction) |

| | | | | | |
|---|---|---|---|---|---|
| | | | | $\bar{x}$ = 4.11, sd = 4.57 (P28 solo)<br>$\bar{x}$ = 54.11, sd = 77.66 (P28 interaction) | |
| Fig. 2 | P20 USV counts x session per context (cohort as random effect) | lmm (MF, FF, MM) | p = 0.01 (MF main effect of session type)*<br>p = 0.056 (FF main effect of session type)<br>p = 0.004 (MM main effect of session type)** | $\bar{x}$ = 2.62, sd = 3.89 (MF solo)<br>$\bar{x}$ = 24.5, sd = 21.37 (MF interaction)<br>$\bar{x}$ = 0, sd = 0(FF solo)<br>$\bar{x}$ = 81.25, sd = 112.87 (FF interaction)<br>$\bar{x}$ = 0, sd = 0 (MM solo)<br>$\bar{x}$ = 30.12, sd = 28.42 (MM interaction) | Excluded cases with more than 11 solo session USVs from the interacting animals combined |
| Fig. 3A (top panel) | MF time, in seconds, spent x non-vocal behavior category x age (cohort, mouse identity, dyad identity as random effects) | lmm | p < 0.001 (interaction effect between non-vocal behavior category x age)**<br>p = 0.96 (social interaction P12 vs P16)<br>p = 0.99 (social interaction P12 vs P20)<br>p = 0.99 (social interaction P12 vs P24)<br>p = 0.52 (social interaction P12 vs P28)<br>p = 0.99 (social interaction P16 vs P20)<br>p = 0.99 (social interaction P16 vs P24)<br>p = 0.90 (social interaction P16 vs P28)<br>p = 0.99 (social interaction P20 vs P24) | $\bar{x}$ = 0.14, sd = 0.23 (social interaction P12)<br>$\bar{x}$ = 0.20, sd = 0.19 (social interaction P16)<br>$\bar{x}$ = 0.16, sd = 0.09 (social interaction P20)<br>$\bar{x}$ = 0.17, sd = 0.08 (social interaction P24)<br>$\bar{x}$ = 0.29, sd = 0.07 (social interaction P28)<br>$\bar{x}$ = 0.34, sd = 0.40 (huddling P12)<br>$\bar{x}$ = 0.63, sd = 0.20 (huddling P16)<br>$\bar{x}$ = 0.23, sd = 0.23 (huddling P20)<br>$\bar{x}$ = 0.03, sd = 0.04 (huddling P24) | |

| | | | | |
|---|---|---|---|---|
| | | | p = 0.68 (social interaction P20 vs P28)<br>p = 0.77 (social interaction P24 vs P28)<br>p = 0.03 (huddling P12 vs P16)*<br>p = 0.76 (huddling P12 vs P20)<br>p = 0.02 (huddling P12 vs P24)*<br>p = 0.03 (huddling P12 vs P28)*<br>p < 0.001 (huddling P16 vs P20)**<br>p < 0.001 (huddling P16 vs P24)**<br>p < 0.001 (huddling P16 vs P28)**<br>p = 0.31 (huddling P20 vs P24)<br>p = 0.38 (huddling P20 vs P28)<br>p = 0.99 (huddling P24 vs P28)<br>p = 0.004 (not interacting P12 vs P16)**<br>p = 0.88 (not interacting P12 vs P20)<br>p = 0.05 (not interacting P12 vs P24)<br>p = 0.62 (not interacting P12 vs P28)<br>p < 0.001 (not interacting P16 vs P20)**<br>p < 0.001 (not interacting P16 vs P24)**<br>p < 0.001 (not interacting P16 vs P28)**<br>p = 0.38 (not interacting P20 vs P24) | $\bar{x}$ = 0.05, sd = 0.04 (huddling P28)<br>$\bar{x}$ = 0.52, sd = 0.51 (not interacting P12)<br>$\bar{x}$ = 0.17, sd = 0.09 (not interacting P16)<br>$\bar{x}$ = 0.61, sd = 0.25 (not interacting P20)<br>$\bar{x}$ = 0.79, sd = 0.09 (not interacting P24)<br>$\bar{x}$ = 0.66, sd = 0.09 (not interacting P28) |

| | | | | |
|---|---|---|---|---|
| | | | p = 0.99 (not interacting P20 vs P28)<br>p = 0.69 (not interacting P24 vs P28) | |
| Fig. 3A (middle panel) | FF time, in seconds, spent x non-vocal behavior category x age (cohort, mouse identity, dyad identity as random effects) | lmm | $p < 0.001$ (interaction effect between non-vocal behavior category x age)**<br>p = 0.81 (social interaction P12 vs P16)<br>p = 0.84 (social interaction P12 vs P20)<br>p = 0.58 (social interaction P12 vs P24)<br>p = 0.45 (social interaction P12 vs P28)<br>p = 1 (social interaction P16 vs P20)<br>p = 1 (social interaction P16 vs P24)<br>p = 0.98 (social interaction P16 vs P28)<br>p = 0.99 (social interaction P20 vs P24)<br>p = 0.97 (social interaction P20 vs P28)<br>p = 1 (social interaction P24 vs P28)<br>p = 0.002 (huddling P12 vs P16)**<br>p = 0.99 (huddling P12 vs P20)<br>p = 0.06 (huddling P12 vs P24)<br>p = 0.04 (huddling P12 vs P28)* | $\bar{x}$ = 0.07, sd = 0.10 (social interaction P12)<br>$\bar{x}$ = 0.17, sd = 0.09 (social interaction P16)<br>$\bar{x}$ = 0.17, sd = 0.06 (social interaction P20)<br>$\bar{x}$ = 0.21, sd = 0.09 (social interaction P24)<br>$\bar{x}$ = 0.23, sd = 0.09 (social interaction P28)<br>$\bar{x}$ = 0.30, sd = 0.38 (huddling P12)<br>$\bar{x}$ = 0.66, sd = 0.20 (huddling P16)<br>$\bar{x}$ = 0.34, sd = 0.20 (huddling P20)<br>$\bar{x}$ = 0.05, sd = 0.04 (huddling P24)<br>$\bar{x}$ = 0.04, sd = 0.03 (huddling P28)<br>$\bar{x}$ = 0.63, sd = 0.46 (not interacting P12)<br>$\bar{x}$ = 0.16, sd = 0.17 (not interacting P16)<br>$\bar{x}$ = 0.49, sd = 0.19 (not interacting P20)<br>$\bar{x}$ = 0.74, sd = 0.10 (not interacting P24) |

| | | | | |
|---|---|---|---|---|
| | | | p = 0.007 (huddling P16 vs P20)** p < 0.001 (huddling P16 vs P24)** p < 0.001 (huddling P16 vs P28)** p = 0.02 (huddling P20 vs P24)* p = 0.01 (huddling P20 vs P28)* p=0.99 (huddling P24 vs P28) p < 0.001 (not interacting P12 vs P16)** p = 0.6 (not interacting P12 vs P20) p = 0.75 (not interacting P12 vs P24) p = 0.78 (not interacting P12 vs P28) p = 0.005 (not interacting P16 vs P20)** p < 0.001 (not interacting P16 vs P24)** p < 0.001 (not interacting P16 vs P28)** p = 0.07 (not interacting P20 vs P24) p = 0.08 (not interacting P20 vs P28) p = 1 (not interacting P24 vs P28) | $\bar{x}$ = 0.73, sd = 0.11 (not interacting P28) |
| Fig. 3A (bottom panel) | MM time, in seconds, spent x non-vocal behavior category x age (cohort, mouse identity, dyad identity as | lmm | p < 0.001 (interaction effect between non-vocal behavior category x age)** p = 0.90 (social interaction P12 vs P16) | $\bar{x}$ = 0.09, sd = 0.16 (social interaction P12) $\bar{x}$ = 0.14, sd = 0.06 (social interaction P16) |

| | | | | |
|---|---|---|---|---|
| | random effects) | | p = 0.14 (social interaction P12 vs P20)<br>p = 0.21 (social interaction P12 vs P24)<br>p = 0.60 (social interaction P12 vs P28)<br>p = 0.60 (social interaction P16 vs P20)<br>p = 0.71 (social interaction P16 vs P24)<br>p = 0.98 (social interaction P16 vs P28)<br>p = 1 (social interaction P20 vs P24)<br>p = 0.92 (social interaction P20 vs P28)<br>p = 0.97 (social interaction P24 vs P28)<br>p < 0.001 (huddling P12 vs P16)**<br>p = 0.98 (huddling P12 vs P20)<br>p = 0.87 (huddling P12 vs P24)<br>p = 0.82 (huddling P12 vs P28)<br>p < 0.001 (huddling P16 vs P20)**<br>p < 0.001 (huddling P16 vs P24)**<br>p < 0.001 (huddling P16 vs P28)**<br>p = 0.57 (huddling P20 vs P24)<br>p = 0.51 (huddling P20 vs P28)<br>p = 1 (huddling P24 vs P28)<br>p < 0.001 (not interacting P12 vs P16)** | $\bar{x}$ = 0.24, sd = 0.11 (social interaction P20)<br>$\bar{x}$ = 0.23, sd = 0.08 (social interaction P24)<br>$\bar{x}$ = 0.18, sd = 0.04 (social interaction P28)<br>$\bar{x}$ = 0.09, sd = 0.17 (huddling P12)<br>$\bar{x}$ = 0.63, sd = 0.24 (huddling P16)<br>$\bar{x}$ = 0.13, sd = 0.1 (huddling P20)<br>$\bar{x}$ = 0.03, sd = 0.02 (huddling P24)<br>$\bar{x}$ = 0.02, sd = 0.02 (huddling P28)<br>$\bar{x}$ = 0.82, sd = 0.33 (not interacting P12)<br>$\bar{x}$ = 0.23, sd = 0.21 (not interacting P16)<br>$\bar{x}$ = 0.63, sd = 0.13 (not interacting P20)<br>$\bar{x}$ = 0.74, sd = 0.77 (not interacting P24)<br>$\bar{x}$ = 0.80, sd = 0.03 (not interacting P28) |

| | | | | |
|---|---|---|---|---|
| | | | p = 0.04 (not interacting P12 vs P20)*<br>p = 0.76 (not interacting P12 vs P24)<br>p = 1 (not interacting P12 vs P28)<br>p < 0.001 (not interacting P16 vs P20)**<br>p < 0.001 (not interacting P16 vs P24)**<br>p < 0.001 (not interacting P16 vs P28)**<br>p = 0.45 (not interacting P20 vs P24)<br>p = 0.12 (not interacting P20 vs P28)<br>p = 0.94 (not interacting P24 vs P28) | |
| Fig. 3B (top panel) | MF mean social interaction bout length x age (cohort, mouse identity, dyad identity as random effects) | glmm-nb | p < 0.001 (main effect of age)**<br>p = 0.06 (P12 vs P16)<br>p < 0.001 (P12 vs P20)**<br>p < 0.001 (P12 vs P24)**<br>p < 0.001 (P12 vs P28)**<br>p = 0.04 (P16 vs P20)*<br>p = 0.002 (P16 vs P24)**<br>p = 0.98 (P20 vs P24)<br>p = 1 (P20 vs P28)<br>p = 1 (P24 vs P28) | $\bar{x}$ = 42.75, sd = 40.51 (P12)<br>$\bar{x}$ = 19.62, sd = 19.53 (P16)<br>$\bar{x}$ = 7.58, sd = 4.97 (P20)<br>$\bar{x}$ = 7.36, sd = 4.3 (P24)<br>$\bar{x}$ = 5.27, sd = 2.57 (P28) |
| Fig. 3B (middle panel) | FF mean social interaction bout length x | glmm-nb | p < 0.001 (main effect of age)**<br>p = 0.48 (P12 vs P16) | $\bar{x}$ = 9.7, sd = 2.18 (P12)<br>$\bar{x}$ = 13.47, sd = 5.38 (P16) |

| | | | | |
|---|---|---|---|---|
| | age (cohort, mouse identity, dyad identity as random effects) | | p = 0.58 (P12 vs P20)<br>p = 0.25 (P12 vs P24)<br>p < 0.001 (P12 vs P28)**<br>p = 0.004 (P16 vs P20)*<br>p < 0.001 (P16 vs P4)**<br>p < 0.001 (P16 vs P8)**<br>p = 0.99 (P20 vs P24)<br>p = 0.03 (P20 vs P28)*<br>p = 0.1 (P24 vs P28) | $\bar{x}$ = 7.28, sd = 2.34 (P20)<br>$\bar{x}$ = 6.66, sd = 3.97 (P24)<br>$\bar{x}$ = 3.92, sd = 1.37 (P28) |
| Fig. 3B (bottom panel) | MM mean social interaction bout length x age (cohort, mouse identity, dyad identity as random effects) | glmm-nb | p < 0.001 (main effect of age)**<br>p < 0.001 (P12 vs P16)**<br>p < 0.001 (P12 vs P20)**<br>p < 0.001 (P12 vs P24)**<br>p < 0.001 (P12 vs P28)**<br>p = 0.79 (P16 vs P20)<br>p = 0.02 (P16 vs P24) *<br>p < 0.001 (P16 vs P28)**<br>p = 0.31 (P20 vs P24)<br>p = 0.04 (P20 vs P28)*<br>p = 0.84 (P24 vs P28) | $\bar{x}$ = 28.2, sd = 26.10 (P12)<br>$\bar{x}$ = 8.99, sd = 5 (P16)<br>$\bar{x}$ = 6.95, sd = 4.63 (P20)<br>$\bar{x}$ = 4.39, sd = 1.53 (P24)<br>$\bar{x}$ = 2.79, sd = 0.39 (P28) |
| Fig. 3C (top panel) | MF number of social interaction bouts x age (cohort, mouse identity, dyad identity as random effects) | lmm | p < 0.001 (main effect of age)**<br>p = 0.99 (P12 vs P16)<br>p = 0.56 (P12 vs P20)<br>p = 0.09 (P12 vs P24) | $\bar{x}$ = 3.4, sd = 0.18 (P12)<br>$\bar{x}$ = 8.5, sd = 6.9 (P16)<br>$\bar{x}$ = 16.5, sd = 10.49 (P20)<br>$\bar{x}$ = 21.1, sd = 17.57 (P24) |

| | | | | |
|---|---|---|---|---|
| | | | p < 0.001 (P12 vs P28)**<br>p = 0.12 (P16 vs P20)<br>p = 0.004 (P16 vs P24)**<br>p < 0.001 (P16 vs P28)**<br>p = 0.61 (P20 vs P24)<br>p < 0.001 (P20 vs P28)**<br>p < 0.001 (P24 vs P28)* | x̅ = 37, sd = 9.99 (P28) |
| Fig. 3C (middle panel) | FF number of social interaction bouts x age (cohort, mouse identity, dyad identity as random effects) | lmm | p < 0.001 (main effect of age)**<br>p = 0.99 (P12 vs P16)<br>p = 0.98 (P12 vs P20)<br>p = 0.35 (P12 vs P24)<br>p = 0.01 (P12 vs P28)*<br>p = 0.71 (P16 vs P20)<br>p = 0.03 (P16 vs P24)*<br>p < 0.001 (P16 vs P28)**<br>p = 0.4 (P20 vs P24)<br>p = 0.002 (P20 vs P28)**<br>p = 0.21 (P24 vs P28) | x̅ = 1.75, sd = 5.73 (P12)<br>x̅ = 8.78, sd = 6.12 (P16)<br>x̅ = 15.33, sd = 8.14 (P20)<br>x̅ = 24.78, sd = 17.18 (P24)<br>x̅ = 36.33, sd = 9.86 (P28) |
| Fig. 3C (bottom panel) | MM number of social interaction bouts x age (cohort, mouse identity, dyad identity as random effects) | lmm | p < 0.001 (main effect of age)**<br>p = 0.51 (P12 vs P16)<br>p = 0.002 (P12 vs P20)**<br>p < 0.001 (P12 vs P24)**<br>p < 0.001 (P12 vs P28)**<br>p = 0.003 (P16 vs P20)* | x̅ = 3.2, sd = 1.64 (P12)<br>x̅ = 11.5, sd = 6.02 (P16)<br>x̅ = 26.7, sd = 11.31 (P20)<br>x̅ = 32.7, sd = 7.50 (P24)<br>x̅ = 39.89, sd = 8.25 (P28) |

| | | | | |
|---|---|---|---|---|
| | | | p < 0.001 (P16 vs P24)**<br>p < 0.001 (P16 vs P28)**<br>p = 0.51 (P20 vs P24)<br>p = 0.01 (P20 vs P28)*<br>p = 0.39 (P24 vs P28) | |
| Fig. 5 | USVs per second behavior per context x age (cohort, mouse identity, dyad identity as random effects) | lmm (MF, FF, MM) | p < 0.001 (MF interaction effect age vs non-vocal behavior category)**<br>p = 0.56 (MF P24 social interaction vs huddling)<br>p = 0.63 (MF P24 social interaction vs not interacting)<br>p = 0.99 (MF P24 huddling vs not interacting)<br>p < 0.001 (MF P28 social interaction vs huddling)**<br>p < 0.001 (MF P28 social interaction vs not interacting)**<br>p = 0.38 (MF P28 huddling vs not interacting)<br>p = 0.03 (FF interaction effect age vs non-vocal behavior category)**<br>p = 0.28 (FF P24 social interaction vs huddling)<br>p = 0.26 (FF P24 social interaction vs not interacting)<br>p = 0.99 (FF P24 huddling vs not interacting)<br>p = 0.82 (FF P28 social interaction vs huddling) | $\bar{x}$ = 0.23, sd = 0.18 (MF P24 social interaction)<br>$\bar{x}$ = 0.35, sd = 0.67 (FF P24 social interaction)<br>$\bar{x}$ = 0.2, sd = 0.23 (MM P24 social interaction)<br>$\bar{x}$ = 0, sd = 0 (MF P24 huddling)<br>$\bar{x}$ = 0.01, sd = 0.03 (FF P24 huddling)<br>$\bar{x}$ = 0, sd = 0 (MM P24 huddling)<br>$\bar{x}$ = 0.03, sd = 0.05 (MF P24 not interacting)<br>$\bar{x}$ = 0.003, sd = 0.002 (FF P24 not interacting)<br>$\bar{x}$ = 0.03, sd = 0.05 (MM P24 not interacting)<br>$\bar{x}$ = 1.26, sd = 1.1 (MF P28 social interaction)<br>$\bar{x}$ = 0.11, sd = 0.16 (FF P28 social interaction)<br>$\bar{x}$ = 0.6, sd = 0.5 (MM P28 social interaction)<br>$\bar{x}$ = 0.35, sd = 1.1 (MF P28 huddling)<br>$\bar{x}$ = 0, sd = 0 (FF P28 huddling) |

| | | | | |
|---|---|---|---|---|
| | | | p = 0.85 (FF P28 social interaction vs not interacting)<br>p = 1 (FF P28 huddling vs not interacting)<br>p = 0.02 (MM interaction effect age vs non-vocal behavior category)**<br>p = 0.23 (MM P24 social interaction vs huddling)<br>p = 0.34 (MM P24 social interaction vs not interacting)<br>p = 0.97 (MM P24 huddling vs not interacting)<br>p = 0.002 (MM P28 social interaction vs huddling)**<br>p = 0.003 (MM P28 social interaction vs not interacting)**<br>p = 1 (MM P28 huddling vs not interacting) | $\bar{x}$ = 0.03, sd = 0.08 (MM P28 huddling)<br>$\bar{x}$ = 0.05, sd = 0.08 (MF P28 not interacting)<br>$\bar{x}$ = 0.01, sd = 0.02 (FF P28 not interacting)<br>$\bar{x}$ = 0.04, sd = 0.04 (MM P28 not interacting) |
| Fig. 6B (top panel) | MF proportion of time spent in USVs vs social interaction (behavior category) per age x time bin (dyad identity as random effect) | lmm | p < 0.001 (P24 interaction effect behavior category vs time bin)**<br>p < 0.001 (P24 time bin #1 proportion of time spent in USVs vs social interaction)**<br>p = 0.6 (P24 time bin #2 proportion of time spent in USVs vs social interaction)<br>p = 0.29 (P24 time bin #3 proportion of time spent in USVs vs social interaction)<br>p = 0.82 (P24 time bin #4 proportion of time spent in USVs | $\bar{x}$ = 0.95, sd = 0.08 (P24 time bin #1 proportion of USVs)<br>$\bar{x}$ = 0.51, sd = 0.24 (P24 time bin #1 proportion of social interaction)<br>$\bar{x}$ = 0.03, sd = 0.05 (P24 time bin #2 proportion of USVs)<br>$\bar{x}$ = 0.12, sd = 0.09 (P24 time bin #2 proportion of social interaction)<br>$\bar{x}$ = 0, sd = 0 (P24 time bin #3 proportion of USVs)<br>$\bar{x}$ = 0.11, sd = 0.09 (P24 time bin |

| | | | | |
|---|---|---|---|---|
| | | | vs social interaction)<br>p = 0.86 (P24 time bin #5 proportion of time spent in USVs vs social interaction)<br>p = 0.63 (P24 time bin #6 proportion of time spent in USVs vs social interaction)<br>p < 0.001 (P28 interaction effect behavior category vs time bin)**<br>p < 0.001 (P28 time bin #1 proportion of time spent in USVs vs social interaction)**<br>p = 0.03 (P28 time bin #2 proportion of time spent in USVs vs social interaction)*<br>p = 0.58 (P28 time bin #3 proportion of time spent in USVs vs social interaction)<br>p = 0.91 (P28 time bin #4 proportion of time spent in USVs vs social interaction)<br>p = 0.44 (P28 time bin #5 proportion of time spent in USVs vs social interaction)<br>p = 0.79 (P28 time bin #6 proportion of time spent in USVs vs social interaction)<br>p = 0.96 (adult interaction effect behavior category vs time bin) | #3 proportion of social interaction)<br>$\bar{x}$ = 0, sd = 0 (P24 time bin #4 proportion of USVs)<br>$\bar{x}$ = 0.08, sd = 0.06 (P24 time bin #4 proportion of social interaction)<br>$\bar{x}$ = 0.02, sd = 0.06 (P24 time bin #5 proportion of USVs)<br>$\bar{x}$ = 0.09, sd = 0.09 (P24 time bin #5 proportion of social interaction)<br>$\bar{x}$ = 0, sd = 0 (P24 time bin #6 proportion of USVs)<br>$\bar{x}$ = 0.09, sd = 0.1 (P24 time bin #6 proportion of social interaction)<br>$\bar{x}$ = 0.84, sd = 0.2 (P28 time bin #1 proportion of USVs)<br>$\bar{x}$ = 0.34, sd = 0.14 (P28 time bin #1 proportion of social interaction)<br>$\bar{x}$ = 0.09, sd = 0.13 (P28 time bin #2 proportion of USVs)<br>$\bar{x}$ = 0.24, sd = 0.14 (P28 time bin #2 proportion of social interaction)<br>$\bar{x}$ = 0.03, sd = 0.07 (P28 time bin #3 proportion of USVs)<br>$\bar{x}$ = 0.13, sd = 0.06 (P28 time bin #3 proportion of social interaction) |

| | | | | |
|---|---|---|---|---|
| | | | | $\bar{x}$ = 0.03, sd = 0.05 (P28 time bin #4 proportion of USVs)<br>$\bar{x}$ = 0.1, sd = 0.05 (P28 time bin #4 proportion of social interaction)<br>$\bar{x}$ = 0, sd = 0.01 (P28 time bin #5 proportion of USVs)<br>$\bar{x}$ = 0.11, sd = 0.05 (P28 time bin #5 proportion of social interaction)<br>$\bar{x}$ = 0.01, sd = 0.03 (P28 time bin #6 proportion of USVs)<br>$\bar{x}$ = 0.09, sd = 0.04 (P28 time bin #6 proportion of social interaction)<br>$\bar{x}$ = 0.15, sd = 0.2 (adult time bin #1 proportion of USVs)<br>$\bar{x}$ = 0.15, sd = 0.16 (adult time bin #1 proportion of social interaction)<br>$\bar{x}$ = 0.21, sd = 0.25 (adult time bin #2 proportion of USVs)<br>$\bar{x}$ = 0.15, sd = 0.16 (adult time bin #2 proportion of social interaction)<br>$\bar{x}$ = 0.13, sd = 0.12 (adult time bin #3 proportion of USVs)<br>$\bar{x}$ = 0.15, sd = 0.16 (adult time bin #3 proportion of social interaction) |

| | | | | |
|---|---|---|---|---|
| | | | | $\bar{x}$ = 0.14, sd = 0.17 (adult time bin #4 proportion of USVs)<br><br>$\bar{x}$ = 0.17, sd = 0.2 (adult time bin #4 proportion of social interaction)<br>$\bar{x}$ = 0.19, sd = 0.23 (adult time bin #5 proportion of USVs)<br>$\bar{x}$ = 0.17, sd = 0.17 (adult time bin #5 proportion of social interaction)<br>$\bar{x}$ = 0.18, sd = 0.23 (adult time bin #6 proportion of USVs)<br>$\bar{x}$ = 0.2, sd = 0.29 (adult time bin #6 proportion of social interaction) |
| Fig. 6B (middle panel) | FF proportion of time spent in USVs vs social interaction (behavior category) per age x time bin (dyad identity as random effect) | lmm | p = 0.01 (P24 interaction effect behavior category vs time bin)*<br>p = 0.03 (P24 time bin #1 proportion of time spent in USVs vs social interaction)*<br>p = 1 (P24 time bin #2 proportion of time spent in USVs vs social interaction)<br>p = 1 (P24 time bin #3 proportion of time spent in USVs vs social interaction)<br>p = 1 (P24 time bin #4 proportion of time spent in USVs vs social interaction) | $\bar{x}$ = 0.8, sd = 0.45 (P24 time bin #1 proportion of USVs)<br>$\bar{x}$ = 0.37, sd = 0.23 (P24 time bin #1 proportion of social interaction)<br>$\bar{x}$ = 0, sd = 0 (P24 time bin #2 proportion of USVs)<br>$\bar{x}$ = 0.1, sd = 0.05 (P24 time bin #2 proportion of social interaction)<br>$\bar{x}$ = 0.07, sd = 0.15 (P24 time bin #3 proportion of USVs)<br>$\bar{x}$ = 0.14, sd = 0.18 (P24 time bin #3 proportion of social interaction) |

| | | | | |
|---|---|---|---|---|
| | | | p = 0.97 (P24 time bin #5 proportion of time spent in USVs vs social interaction)<br>p = 1 (P24 time bin #6 proportion of time spent in USVs vs social interaction)<br>p = 0.009 (P28 interaction effect behavior category vs time bin)**<br>p = 0.03 (P28 time bin #1 proportion of time spent in USVs vs social interaction)*<br>p = 1 (P28 time bin #2 proportion of time spent in USVs vs social interaction)<br>p = 1 (P28 time bin #3 proportion of time spent in USVs vs social interaction)<br>p = 1 (P28 time bin #4 proportion of time spent in USVs vs social interaction)<br>p = 1 (P28 time bin #5 proportion of time spent in USVs vs social interaction)<br>p = 1 (P28 time bin #6 proportion of time spent in USVs vs social interaction)<br>p = 0.001 (adult interaction effect behavior category vs time bin)**<br>p = 0.01 (adult time bin #1 proportion of time spent in USVs | $\bar{x}$ = 0.13, sd = 0.3 (P24 time bin #4 proportion of USVs)<br>$\bar{x}$ = 0.16, sd = 0.13 (P24 time bin #4 proportion of social interaction)<br>$\bar{x}$ = 0, sd = 0 (P24 time bin #5 proportion of USVs)<br>$\bar{x}$ = 0.16, sd = 0.09 (P24 time bin #5 proportion of social interaction)<br>$\bar{x}$ = 0, sd = 0 (P24 time bin #6 proportion of USVs)<br>$\bar{x}$ = 0.08, sd = 0.06 (P24 time bin #6 proportion of social interaction)<br>$\bar{x}$ = 0.7, sd = 0.48 (P28 time bin #1 proportion of USVs)<br>$\bar{x}$ = 0.3, sd = 0.03 (P28 time bin #1 proportion of social interaction)<br>$\bar{x}$ = 0.15, sd = 0.38 (P28 time bin #2 proportion of USVs)<br>$\bar{x}$ = 0.12, sd = 0.04 (P28 time bin #2 proportion of social interaction)<br>$\bar{x}$ = 0, sd = 0 (P28 time bin #3 proportion of USVs)<br>$\bar{x}$ = 0.15, sd = 0.04 (P28 time bin #3 proportion of social interaction)<br>$\bar{x}$ = 0, sd = 0.01 (P28 time bin #4 |

| | | | | |
|---|---|---|---|---|
| | | | vs social interaction)*<br>p = 1 (adult time bin #2 proportion of time spent in USVs vs social interaction)<br>p = 1 (adult time bin #3 proportion of time spent in USVs vs social interaction)<br>p = 1 (adult time bin #4 proportion of time spent in USVs vs social interaction)<br>p = 0.98 (adult time bin #5 proportion of time spent in USVs vs social interaction)<br>p = 0.71 (adult time bin #6 proportion of time spent in USVs vs social interaction) | proportion of USVs)<br>$\overline{x}$ = 0.12, sd = 0.05 (P28 time bin #4 proportion of social interaction)<br>$\overline{x}$ = 0, sd = 0 (P28 time bin #5 proportion of USVs)<br>$\overline{x}$ = 0.13, sd = 0.05 (P28 time bin #5 proportion of social interaction)<br>$\overline{x}$ = 0.15, sd = 0.37 (P28 time bin #6 proportion of USVs)<br>$\overline{x}$ = 0.18, sd = 0.09 (P28 time bin #6 proportion of social interaction)<br>$\overline{x}$ = 0.68, sd = 0.35 (adult time bin #1 proportion of USVs)<br>$\overline{x}$ = 0.51, sd = 0.22 (adult time bin #1 proportion of social interaction)<br>$\overline{x}$ = 0.12, sd = 0.2 (adult time bin #2 proportion of USVs)<br>$\overline{x}$ = 0.12, sd = 0.2 (adult time bin #2 proportion of social interaction)<br>$\overline{x}$ = 0.07, sd = 0.12 (adult time bin #3 proportion of USVs)<br>$\overline{x}$ = 0.07, sd = 0.1 (adult time bin #3 proportion of social interaction)<br>$\overline{x}$ = 0.05, sd = 0.13 (adult time bin #4 proportion of USVs) |

| | | | | |
|---|---|---|---|---|
| | | | | $\bar{x}$ = 0.07, sd = 0.12 (adult time bin #4 proportion of social interaction)<br>$\bar{x}$ = 0.06, sd = 0.11 (adult time bin #5 proportion of USVs)<br>$\bar{x}$ = 0.12, sd = 0.16 (adult time bin #5 proportion of social interaction)<br>$\bar{x}$ = 0.03, sd = 0.09 (adult time bin #6 proportion of USVs)<br>$\bar{x}$ = 0.12, sd = 0.16 (adult time bin #6 proportion of social interaction) |
| Fig. 6B (bottom panel) | MM proportion of time spent in USVs vs social interaction (behavior category) per age x time bin (dyad identity as random effect) | lmm | $p < 0.001$ (P24 interaction effect behavior category vs time bin)**<br>$p < 0.001$ (P24 time bin #1 proportion of time spent in USVs vs social interaction)**<br>$p < 0.001$ (P24 time bin #2 proportion of time spent in USVs vs social interaction)**<br>$p < 0.001$ (P24 time bin #3 proportion of time spent in USVs vs social interaction)**<br>$p = 0.003$ (P24 time bin #4 proportion of time spent in USVs vs social interaction)**<br>$p < 0.001$ (P24 time bin #5 proportion of time spent in USVs | $\bar{x}$ = 0.96, sd = 0.06 (P24 time bin #1 proportion of USVs)<br>$\bar{x}$ = 0.31, sd = 0.09 (P24 time bin #1 proportion of social interaction)<br>$\bar{x}$ = 0.02, sd = 0.05 (P24 time bin #2 proportion of USVs)<br>$\bar{x}$ = 0.19, sd = 0.06 (P24 time bin #2 proportion of social interaction)<br>$\bar{x}$ = 0.01, sd = 0.03 (P24 time bin #3 proportion of USVs)<br>$\bar{x}$ = 0.16, sd = 0.06 (P24 time bin #3 proportion of social interaction)<br><br>$\bar{x}$ = 0, sd = 0 (P24 time bin #4 |

| | | | | |
|---|---|---|---|---|
| | | | vs social interaction)**<br>p = 0.007 (P24 time bin #6 proportion of time spent in USVs vs social interaction)**<br>p < 0.001 (P28 interaction effect behavior category vs time bin)**<br>p < 0.001 (P28 time bin #1 proportion of time spent in USVs vs social interaction)**<br>p = 0.039 (P28 time bin #2 proportion of time spent in USVs vs social interaction)*<br>p = 0.08 (P28 time bin #3 proportion of time spent in USVs vs social interaction)<br>p = 0.009 (P28 time bin #4 proportion of time spent in USVs vs social interaction)**<br>p = 0.002 (P28 time bin #5 proportion of time spent in USVs vs social interaction)**<br>p = 0.14 (P28 time bin #6 proportion of time spent in USVs vs social interaction)<br>p = 0.54 (adult interaction effect behavior category vs time bin) | proportion of USVs)<br>$\bar{x}$ = 0.11, sd = 0.06 (P24 time bin #4 proportion of social interaction)<br>$\bar{x}$ = 0, sd = 0 (P24 time bin #5 proportion of USVs)<br>$\bar{x}$ = 0.13, sd = 0.05 (P24 time bin #5 proportion of social interaction)<br>$\bar{x}$ = 0, sd = 0 (P24 time bin #6 proportion of USVs)<br>$\bar{x}$ = 0.1, sd = 0.1 (P24 time bin #6 proportion of social interaction)<br>$\bar{x}$ = 0.97, sd = 0.03 (P28 time bin #1 proportion of USVs)<br>$\bar{x}$ = 0.39, sd = 0.12 (P28 time bin #1 proportion of social interaction)<br>$\bar{x}$ = 0.02, sd = 0.02 (P28 time bin #2 proportion of USVs)<br>$\bar{x}$ = 0.13, sd = 0.13 (P28 time bin #2 proportion of social interaction)<br>$\bar{x}$ = 0, sd = 0 (P28 time bin #3 proportion of USVs)<br>$\bar{x}$ = 0.11, sd = 0.05 (P28 time bin #3 proportion of social interaction)<br>$\bar{x}$ = 0, sd = 0.01 (P28 time bin #4 proportion of USVs) |

| | | | | |
|---|---|---|---|---|
| | | | | $\bar{x}$ = 0.13, sd = 0.06 (P28 time bin #4 proportion of social interaction)<br>$\bar{x}$ = 0, sd = 0 (P28 time bin #5 proportion of USVs)<br>$\bar{x}$ = 0.14, sd = 0.07 (P28 time bin #5 proportion of social interaction)<br>$\bar{x}$ = 0, sd = 0 (P28 time bin #6 proportion of USVs)<br>$\bar{x}$ = 0.1, sd = 0.04 (P28 time bin #6 proportion of social interaction)<br>$\bar{x}$ = 0.47, sd = 0.29 (adult time bin #1 proportion of USVs)<br>$\bar{x}$ = 0.58, sd = 0.23 (adult time bin #1 proportion of social interaction)<br>$\bar{x}$ = 0.29, sd = 0.36 (adult time bin #2 proportion of USVs)<br>$\bar{x}$ = 0.13, sd = 0.2 (adult time bin #2 proportion of social interaction)<br>$\bar{x}$ = 0.09, sd = 0.07 (adult time bin #3 proportion of USVs)<br>$\bar{x}$ = 0.04, sd = 0.07 (adult time bin #3 proportion of social interaction)<br>$\bar{x}$ = 0.05, sd = 0.07 (adult time bin #4 proportion of USVs) |

| | | | | | |
|---|---|---|---|---|---|
| | | | | $\bar{x}$ = 0.07, sd = 0.1 (adult time bin #4 proportion of social interaction)<br>$\bar{x}$ = 0.04, sd = 0.05 (adult time bin #5 proportion of USVs)<br>$\bar{x}$ = 0.07, sd = 0.07 (adult time bin #5 proportion of social interaction)<br>$\bar{x}$ = 0.06, sd = 0.08 (adult time bin #6 proportion of USVs)<br>$\bar{x}$ = 0.1, sd = 0.12 (adult time bin #6 proportion of social interaction) | |
| Fig. S1 | interaction session USV rate x age per context | lmm | p = 0.05 (MF main effect of age)*<br>p = 0.46 (MF P24 vs P28)<br>p = 0.04 (MF P24 vs adult)*<br>p = 0.53 (MF P28 vs adult)<br>p = 0.002 (FF main effect of age)**<br>p = 0.99 (FF P24 vs P28)<br>p = 0.01 (FF P24 vs adult)*<br>p = 0.01 (FF P28 vs adult)*<br>p = 0.32 (MM main effect of age) | $\bar{x}$ = 38.1, sd = 31.53 (MF P24 USVs)<br>$\bar{x}$ = 259.8, sd = 248.84 (MF P28 USVs)<br>$\bar{x}$ = 431.27, sd = 541.42 (MF adult USVs)<br>$\bar{x}$ = 20.33, sd = 42.69 (FF P24 USVs)<br>$\bar{x}$ = 21.67, sd = 35.51 (FF P28 USVs)<br>$\bar{x}$ = 240.29, sd = 244.86 (FF adult USVs)<br>$\bar{x}$ = 38.1, sd = 37.92 (MM P24 USVs)<br>$\bar{x}$ = 54.11, sd = 77.66 (MM P28 USVs)<br>$\bar{x}$ = 141.2, sd = 262.05 (MM adult USVs) | analyses performed on P24, P28, and adult ages only<br>for adult data, used USV rates from first 10 minutes |

| | | | | |
|---|---|---|---|---|
| Fig. S2 | proportion of USVs (Fig. S2A) and time spent in social interaction (Fig. S2B) x time bin per context and age (filename as random effect) | lmm | p < 0.001 (MF P24 proportion of USVs main effect of time bin)**<br>p < 0.001 (MF P24 proportion of USVs time bin #1 vs #2)**<br>p < 0.001 (MF P24 proportion of USVs time bin #1 vs #3)**<br>p < 0.001 (MF P24 proportion of USVs time bin #1 vs #4)**<br>p < 0.001 (MF P24 proportion of USVs time bin #1 vs #5)**<br>p < 0.001 (MF P24 proportion of USVs time bin #1 vs #6)**<br>p = 0.75 (MF P24 proportion of USVs time bin #2 vs #3)<br>p = 0.75 (MF P24 proportion of USVs time bin #2 vs #4)<br>p = 0.99 (MF P24 proportion of USVs time bin #2 vs #5)<br>p = 0.75 (MF P24 proportion of USVs time bin #2 vs #6)<br>p = 1 (MF P24 proportion of USVs time bin #3 vs #4)<br>p = 0.92 (MF P24 proportion of USVs time bin #3 vs #5)<br>p = 1 (MF P24 proportion of USVs time bin #3 vs #6)<br>p = 0.92 (MF P24 proportion of USVs time bin #4 vs #5)<br>p = 1 (MF P24 proportion of USVs time bin #4 vs #6)<br>p = 0.92 (MF P24 proportion of USVs time bin #5 vs #6)<br>p < 0.001 (MF P28 proportion of USVs | $\bar{x}$ = 0.95, sd = 0.08 (P24 MF time bin #1 proportion of USVs)<br>$\bar{x}$ = 0.51, sd = 0.24 (P24 MF time bin #1 proportion of social interaction)<br>$\bar{x}$ = 0.03, sd = 0.05 (P24 MF time bin #2 proportion of USVs)<br>$\bar{x}$ = 0.12, sd = 0.09 (P24 MF time bin #2 proportion of social interaction)<br>$\bar{x}$ = 0, sd = 0 (P24 MF time bin #3 proportion of USVs)<br>$\bar{x}$ = 0.11, sd = 0.09 (P24 MF time bin #3 proportion of social interaction)<br>$\bar{x}$ = 0, sd = 0 (P24 MF time bin #4 proportion of USVs)<br>$\bar{x}$ = 0.08, sd = 0.06 (P24 MF time bin #4 proportion of social interaction)<br>$\bar{x}$ = 0.02, sd = 0.06 (P24 MF time bin #5 proportion of USVs)<br>$\bar{x}$ = 0.09, sd = 0.09 (P24 MF time bin #5 proportion of social interaction)<br>$\bar{x}$ = 0, sd = 0 (P24 MF time bin #6 |

| | | | | |
|---|---|---|---|---|
| | | | main effect of time bin)**<br>$p < 0.001$ (MF P28 proportion of USVs time bin #1 vs #2)**<br>$p < 0.001$ (MF P28 proportion of USVs time bin #1 vs #3)**<br>$p < 0.001$ (MF P28 proportion of USVs time bin #1 vs #4)**<br>$p < 0.001$ (MF P28 proportion of USVs time bin #1 vs #5)**<br>$p < 0.001$ (MF P28 proportion of USVs time bin #1 vs #6)**<br>$p = 0.85$ (MF P28 proportion of USVs time bin #2 vs #3)<br>$p = 0.83$ (MF P28 proportion of USVs time bin #2 vs #4)<br>$p = 0.49$ (MF P28 proportion of USVs time bin #2 vs #5)<br>$p = 0.57$ (MF P28 proportion of USVs time bin #2 vs #6)<br>$p = 1$ (MF P28 proportion of USVs time bin #3 vs #4)<br>$p = 0.99$ (MF P28 proportion of USVs time bin #3 vs #5)<br>$p = 0.99$ (MF P28 proportion of USVs time bin #3 vs #6)<br>$p = 0.99$ (MF P28 proportion of USVs time bin #4 vs #5)<br>$p = 1$ (MF P28 proportion of USVs time bin #4 vs #6)<br>$p = 1$ (MF P28 proportion of USVs time bin #5 vs #6)<br>$p = 0.9$ (MF adult proportion of USVs main effect of time bin) | proportion of USVs)<br>$\bar{x} = 0.09$, sd = 0.1 (P24 MF time bin #6 proportion of social interaction)<br>$\bar{x} = 0.84$, sd = 0.2 (P28 MF time bin #1 proportion of USVs)<br>$\bar{x} = 0.34$, sd = 0.14 (P28 MF time bin #1 proportion of social interaction)<br>$\bar{x} = 0.09$, sd = 0.13 (P28 MF time bin #2 proportion of USVs)<br>$\bar{x} = 0.24$, sd = 0.14 (P28 MF time bin #2 proportion of social interaction)<br>$\bar{x} = 0.03$, sd = 0.07 (P28 MF time bin #3 proportion of USVs)<br>$\bar{x} = 0.13$, sd = 0.06 (P28 MF time bin #3 proportion of social interaction)<br>$\bar{x} = 0.03$, sd = 0.05 (P28 MF time bin #4 proportion of USVs)<br>$\bar{x} = 0.1$, sd = 0.05 (P28 MF time bin #4 proportion of social interaction)<br>$\bar{x} = 0$, sd = 0.01 (P28 MF time bin #5 proportion of USVs)<br>$\bar{x} = 0.11$, sd = 0.05 (P28 MF time bin #5 |

| | | | | |
|---|---|---|---|---|
| | | | $p < 0.001$ (FF P24 proportion of USVs main effect of time bin)**<br>$p < 0.001$ (FF P24 proportion of USVs time bin #1 vs #2)**<br>$p < 0.001$ (FF P24 proportion of USVs time bin #1 vs #3)**<br>$p = 0.002$ (FF P24 proportion of USVs time bin #1 vs #4)**<br>$p < 0.001$ (FF P24 proportion of USVs time bin #1 vs #5)**<br>$p < 0.001$ (FF P24 proportion of USVs time bin #1 vs #6)**<br>$p = 1$ (FF P24 proportion of USVs time bin #2 vs #3)<br>$p = 0.94$ (FF P24 proportion of USVs time bin #2 vs #4)<br>$p = 1$ (FF P24 proportion of USVs time bin #2 vs #5)<br>$p = 1$ (FF P24 proportion of USVs time bin #2 vs #6)<br>$p = 1$ (FF P24 proportion of USVs time bin #3 vs #4)<br>$p = 1$ (FF P24 proportion of USVs time bin #3 vs #5)<br>$p = 1$ (FF P24 proportion of USVs time bin #3 vs #6)<br>$p = 0.94$ (FF P24 proportion of USVs time bin #4 vs #5)<br>$p = 0.94$ (FF P24 proportion of USVs time bin #4 vs #6)<br>$p = 1$ (FF P24 proportion of USVs time bin #5 vs #6)<br>$p < 0.001$ (FF P28 proportion of USVs | proportion of social interaction)<br>$\bar{x} = 0.01$, sd = 0.03 (P28 MF time bin #6 proportion of USVs)<br>$\bar{x} = 0.09$, sd = 0.04 (P28 MF time bin #6 proportion of social interaction)<br>$\bar{x} = 0.15$, sd = 0.2 (adult MF time bin #1 proportion of USVs)<br>$\bar{x} = 0.15$, sd = 0.16 (adult MF time bin #1 proportion of social interaction)<br>$\bar{x} = 0.21$, sd = 0.25 (adult MF time bin #2 proportion of USVs)<br>$\bar{x} = 0.15$, sd = 0.16 (adult MF time bin #2 proportion of social interaction)<br>$\bar{x} = 0.13$, sd = 0.12 (adult MF time bin #3 proportion of USVs)<br>$\bar{x} = 0.15$, sd = 0.16 (adult MF time bin #3 proportion of social interaction)<br>$\bar{x} = 0.14$, sd = 0.17 (adult MF time bin #4 proportion of USVs)<br>$\bar{x} = 0.17$, sd = 0.2 (adult MF time bin #4 proportion of social interaction) |

| | | | | |
|---|---|---|---|---|
| | | | main effect of time bin)**<br>p = 0.02 (FF P28 proportion of USVs time bin #1 vs #2)*<br>p = 0.001 (FF P28 proportion of USVs time bin #1 vs #3)**<br>p = 0.001 (FF P28 proportion of USVs time bin #1 vs #4)**<br>p = 0.001 (FF P28 proportion of USVs time bin #1 vs #5)**<br>p = 0.02 (FF P28 proportion of USVs time bin #1 vs #6)*<br>p = 0.93 (FF P28 proportion of USVs time bin #2 vs #3)<br>p = 0.94 (FF P28 proportion of USVs time bin #2 vs #4)<br>p = 0.94 (FF P28 proportion of USVs time bin #2 vs #5)<br>p = 1 (FF P28 proportion of USVs time bin #2 vs #6)<br>p = 1 (FF P28 proportion of USVs time bin #3 vs #4)<br>p = 1 (FF P28 proportion of USVs time bin #3 vs #5)<br>p = 0.92 (FF P28 proportion of USVs time bin #3 vs #6)<br>p = 1 (FF P28 proportion of USVs time bin #4 vs #5)<br>p = 0.93 (FF P28 proportion of USVs time bin #4 vs #6)<br>p = 0.92 (FF P28 proportion of USVs time bin #5 vs #6)<br>p < 0.001 (FF adult proportion of USVs main effect of time bin)** | $\bar{x}$ = 0.19, sd = 0.23 (adult MF time bin #5 proportion of USVs)<br>$\bar{x}$ = 0.17, sd = 0.17 (adult MF time bin #5 proportion of social interaction)<br>$\bar{x}$ = 0.18, sd = 0.23 (adult MF time bin #6 proportion of USVs)<br>$\bar{x}$ = 0.2, sd = 0.29 (adult MF time bin #6 proportion of social interaction)<br>$\bar{x}$ = 0.8, sd = 0.45 (P24 FF time bin #1 proportion of USVs)<br>$\bar{x}$ = 0.37, sd = 0.23 (P24 FF time bin #1 proportion of social interaction)<br>$\bar{x}$ = 0, sd = 0 (P24 FF time bin #2 proportion of USVs)<br>$\bar{x}$ = 0.1, sd = 0.05 (P24 FF time bin #2 proportion of social interaction)<br>$\bar{x}$ = 0.07, sd = 0.15 (P24 FF time bin #3 proportion of USVs)<br>$\bar{x}$ = 0.14, sd = 0.18 (P24 FF time bin #3 proportion of social interaction)<br>$\bar{x}$ = 0.13, sd = 0.3 (P24 FF time bin #4 proportion of USVs)<br>$\bar{x}$ = 0.16, sd = 0.13 (P24 FF time |

| | | | | |
|---|---|---|---|---|
| | | | p < 0.001 (FF adult proportion of USVs time bin #1 vs #2)**<br>p < 0.001 (FF adult proportion of USVs time bin #1 vs #3)**<br>p < 0.001 (FF adult proportion of USVs time bin #1 vs #4)**<br>p < 0.001 (FF adult proportion of USVs time bin #1 vs #5)**<br>p < 0.001 (FF adult proportion of USVs time bin #1 vs #6)**<br>p = 0.87 (FF adult proportion of USVs time bin #2 vs #3)<br>p = 0.68 (FF adult proportion of USVs time bin #2 vs #4)<br>p = 0.83 (FF adult proportion of USVs time bin #2 vs #5)<br>p = 0.40 (FF adult proportion of USVs time bin #2 vs #6)<br>p = 1 (FF adult proportion of USVs time bin #3 vs #4)<br>p = 1 (FF adult proportion of USVs time bin #3 vs #5)<br>p = 0.97 (FF adult proportion of USVs time bin #3 vs #6)<br>p = 1 (FF adult proportion of USVs time bin #4 vs #5)<br>p = 1 (FF adult proportion of USVs time bin #4 vs #6)<br>p = 0.98 (FF adult proportion of USVs time bin #5 vs #6)<br>p < 0.001 (MM P24 proportion of USVs main effect of time bin)** | bin #4 proportion of social interaction)<br>$\bar{x}$ = 0, sd = 0 (P24 FF time bin #5 proportion of USVs)<br>$\bar{x}$ = 0.16, sd = 0.09 (P24 FF time bin #5 proportion of social interaction)<br>$\bar{x}$ = 0, sd = 0 (P24 FF time bin #6 proportion of USVs)<br>$\bar{x}$ = 0.08, sd = 0.06 (P24 FF time bin #6 proportion of social interaction)<br>$\bar{x}$ = 0.7, sd = 0.48 (P28 FF time bin #1 proportion of USVs)<br>$\bar{x}$ = 0.3, sd = 0.03 (P28 FF time bin #1 proportion of social interaction)<br>$\bar{x}$ = 0.15, sd = 0.38 (P28 FF time bin #2 proportion of USVs)<br>$\bar{x}$ = 0.12, sd = 0.04 (P28 FF time bin #2 proportion of social interaction)<br>$\bar{x}$ = 0, sd = 0 (P28 FF time bin #3 proportion of USVs)<br>$\bar{x}$ = 0.15, sd = 0.04 (P28 FF time bin #3 proportion of social interaction)<br>$\bar{x}$ = 0, sd = 0.01 (P28 FF time bin #4 proportion of USVs) |

| | | | | |
|---|---|---|---|---|
| | | | $p < 0.001$ (MM P24 proportion of USVs time bin #1 vs #2)**<br>$p < 0.001$ (MM P24 proportion of USVs time bin #1 vs #3)**<br>$p < 0.001$ (MM P24 proportion of USVs time bin #1 vs #4)**<br>$p < 0.001$ (MM P24 proportion of USVs time bin #1 vs #5)**<br>$p < 0.001$ (MM P24 proportion of USVs time bin #1 vs #6)**<br>$p = 1$ (MM P24 proportion of USVs time bin #2 vs #3)<br>$p = 0.67$ (MM P24 proportion of USVs time bin #2 vs #4)<br>$p = 0.67$ (MM P24 proportion of USVs time bin #2 vs #5)<br>$p = 0.67$ (MM P24 proportion of USVs time bin #2 vs #6)<br>$p = 0.93$ (MM P24 proportion of USVs time bin #3 vs #4)<br>$p = 0.93$ (MM P24 proportion of USVs time bin #3 vs #5)<br>$p = 0.93$ (MM P24 proportion of USVs time bin #3 vs #6)<br>$p = 1$ (MM P24 proportion of USVs time bin #4 vs #5)<br>$p = 1$ (MM P24 proportion of USVs time bin #4 vs #6)<br>$p = 1$ (MM P24 proportion of USVs time bin #5 vs #6)<br>$p < 0.001$ (MM P28 proportion of USVs main effect of time bin)** | $\bar{x} = 0.12$, sd = 0.05 (P28 FF time bin #4 proportion of social interaction)<br>$\bar{x} = 0$, sd = 0 (P28 FF time bin #5 proportion of USVs)<br>$\bar{x} = 0.13$, sd = 0.05 (P28 FF time bin #5 proportion of social interaction)<br>$\bar{x} = 0.15$, sd = 0.37 (P28 FF time bin #6 proportion of USVs)<br>$\bar{x} = 0.18$, sd = 0.09 (P28 FF time bin #6 proportion of social interaction)<br>$\bar{x} = 0.68$, sd = 0.35 (adult FF time bin #1 proportion of USVs)<br>$\bar{x} = 0.51$, sd = 0.22 (adult FF time bin #1 proportion of social interaction)<br>$\bar{x} = 0.12$, sd = 0.2 (adult FF time bin #2 proportion of USVs)<br>$\bar{x} = 0.12$, sd = 0.2 (adult FF time bin #2 proportion of social interaction)<br>$\bar{x} = 0.07$, sd = 0.12 (adult FF time bin #3 proportion of USVs)<br>$\bar{x} = 0.07$, sd = 0.1 (adult FF time bin #3 proportion of social interaction) |

| | | | | |
|---|---|---|---|---|
| | | | p < 0.001 (MM P28 proportion of USVs time bin #1 vs #2)**<br>p < 0.001 (MM P28 proportion of USVs time bin #1 vs #3)**<br>p < 0.001 (MM P28 proportion of USVs time bin #1 vs #4)**<br>p < 0.001 (MM P28 proportion of USVs time bin #1 vs #5)**<br>p < 0.001 (MM P28 proportion of USVs time bin #1 vs #6)**<br>p = 0.37 (MM P28 proportion of USVs time bin #2 vs #3)<br>p = 0.49 (MM P28 proportion of USVs time bin #2 vs #4)<br>p = 0.27 (MM P28 proportion of USVs time bin #2 vs #5)<br>p = 0.27 (MM P28 proportion of USVs time bin #2 vs #6)<br>p = 1 (MM P28 proportion of USVs time bin #3 vs #4)<br>p = 1 (MM P28 proportion of USVs time bin #3 vs #5)<br>p = 1 (MM P28 proportion of USVs time bin #3 vs #6)<br>p = 1 (MM P28 proportion of USVs time bin #3 vs #4)<br>p = 1 (MM P28 proportion of USVs time bin #4 vs #5)<br>p = 1 (MM P28 proportion of USVs time bin #4 vs #6)<br>p = 1 (MM P28 proportion of USVs time bin #5 vs #6)<br>p = 0.003 (MM adult proportion of | $\bar{x}$ = 0.05, sd = 0.13 (adult FF time bin #4 proportion of USVs)<br>$\bar{x}$ = 0.07, sd = 0.12 (adult FF time bin #4 proportion of social interaction)<br>$\bar{x}$ = 0.06, sd = 0.11 (adult FF time bin #5 proportion of USVs)<br>$\bar{x}$ = 0.12, sd = 0.16 (adult FF time bin #5 proportion of social interaction)<br>$\bar{x}$ = 0.03, sd = 0.09 (adult FF time bin #6 proportion of USVs)<br>$\bar{x}$ = 0.12, sd = 0.16 (adult FF time bin #6 proportion of social interaction)<br>$\bar{x}$ = 0.96, sd = 0.06 (P24 MM time bin #1 proportion of USVs)<br>$\bar{x}$ = 0.31, sd = 0.09 (P24 MM time bin #1 proportion of social interaction)<br>$\bar{x}$ = 0.02, sd = 0.05 (P24 MM time bin #2 proportion of USVs)<br>$\bar{x}$ = 0.19, sd = 0.06 (P24 MM time bin #2 proportion of social interaction) |

| | | | | |
|---|---|---|---|---|
| | | | USVs main effect of time bin)**<br>p = 0.59 (MM adult proportion of USVs time bin #1 vs #2)<br>p = 0.02 (MM adult proportion of USVs time bin #1 vs #3)*<br>p = 0.01 (MM adult proportion of USVs time bin #1 vs #4)*<br>p = 0.009 (MM adult proportion of USVs time bin #1 vs #5)**<br>p = 0.01 (MM adult proportion of USVs time bin #1 vs #6)*<br>p = 0.48 (MM adult proportion of USVs time bin #2 vs #3)<br>p = 0.31 (MM adult proportion of USVs time bin #2 vs #4)<br>p = 0.26 (MM adult proportion of USVs time bin #2 vs #5)<br>p = 0.33 (MM adult proportion of USVs time bin #2 vs #6)<br>p = 1 (MM adult proportion of USVs time bin #3 vs #4)<br>p = 1 (MM adult proportion of USVs time bin #3 vs #5)<br>p = 1 (MM adult proportion of USVs time bin #3 vs #6)<br>p < 0.001 (MF P24 proportion of social interaction main effect of time bin)**<br>p < 0.001 (MF P24 proportion of social interaction time bin #1 vs #2)**<br>p < 0.001 (MF P24 proportion of social interaction time bin #1 vs #3)** | $\bar{x}$ = 0.01, sd = 0.03 (P24 MM time bin #3 proportion of USVs)<br>$\bar{x}$ = 0.16, sd = 0.06 (P24 MM time bin #3 proportion of social interaction)<br>$\bar{x}$ = 0, sd = 0 (P24 MM time bin #4 proportion of USVs)<br>$\bar{x}$ = 0.11, sd = 0.06 (P24 MM time bin #4 proportion of social interaction)<br>$\bar{x}$ = 0, sd = 0 (P24 MM time bin #5 proportion of USVs)<br>$\bar{x}$ = 0.13, sd = 0.05 (P24 MM time bin #5 proportion of social interaction)<br>$\bar{x}$ = 0, sd = 0 (P24 MM time bin #6 proportion of USVs)<br>$\bar{x}$ = 0.1, sd = 0.1 (P24 MM time bin #6 proportion of social interaction)<br>$\bar{x}$ = 0.97, sd = 0.03 (P28 MM time bin #1 proportion of USVs)<br>$\bar{x}$ = 0.39, sd = 0.12 (P28 MM time bin #1 proportion of social interaction)<br>$\bar{x}$ = 0.02, sd = 0.02 (P28 MM time bin #2 proportion of USVs) |

| | | | | |
|---|---|---|---|---|
| | | | $p < 0.001$ (MF P24 proportion of social interaction time bin #1 vs #4)**<br>$p < 0.001$ (MF P24 proportion of social interaction time bin #1 vs #5)**<br>$p < 0.001$ (MF P24 proportion of social interaction time bin #1 vs #6)**<br>$p = 1$ (MF P24 proportion of social interaction time bin #2 vs #3)<br>$p = 0.98$ (MF P24 proportion of social interaction time bin #2 vs #4)<br>$p = 1$ (MF P24 proportion of social interaction time bin #2 vs #5)<br>$p = 1$ (MF P24 proportion of social interaction time bin #2 vs #6)<br>$p = 0.99$ (MF P24 proportion of social interaction time bin #3 vs #4)<br>$p = 1$ (MF P24 proportion of social interaction time bin #3 vs #5)<br>$p = 1$ (MF P24 proportion of social interaction time bin #3 vs #6)<br>$p = 1$ (MF P24 proportion of social interaction time bin #4 vs #5)<br>$p = 1$ (MF P24 proportion of social interaction time bin #4 vs #6)<br>$p = 1$ (MF P24 proportion of social | $\bar{x} = 0.13$, sd = 0.13 (P28 MM time bin #2 proportion of social interaction)<br>$\bar{x} = 0$, sd = 0 (P28 MM time bin #3 proportion of USVs)<br>$\bar{x} = 0.11$, sd = 0.05 (P28 MM time bin #3 proportion of social interaction)<br>$\bar{x} = 0$, sd = 0.01 (P28 MM time bin #4 proportion of USVs)<br>$\bar{x} = 0.13$, sd = 0.06 (P28 MM time bin #4 proportion of social interaction)<br>$\bar{x} = 0$, sd = 0 (P28 MM time bin #5 proportion of USVs)<br>$\bar{x} = 0.14$, sd = 0.07 (P28 MM time bin #5 proportion of social interaction)<br>$\bar{x} = 0$, sd = 0 (P28 MM time bin #6 proportion of USVs)<br>$\bar{x} = 0.1$, sd = 0.04 (P28 MM time bin #6 proportion of social interaction)<br>$\bar{x} = 0.47$, sd = 0.29 (adult MM time bin #1 proportion of USVs)<br>$\bar{x} = 0.58$, sd = 0.23 (adult MM time bin #1 proportion of social interaction) |

| | | | | |
|---|---|---|---|---|
| | | | interaction time bin #5 vs #6)<br>p < 0.001 (MF P28 proportion of social interaction main effect of time bin)**<br>p = 0.1 (MF P28 proportion of social interaction time bin #1 vs #2)<br>p < 0.001 (MF P28 proportion of social interaction time bin #1 vs #3)**<br>p < 0.001 (MF P28 proportion of social interaction time bin #1 vs #4)**<br>p < 0.001 (MF P28 proportion of social interaction time bin #1 vs #5)**<br>p < 0.001 (MF P28 proportion of social interaction time bin #1 vs #6)**<br>p = 0.07 (MF P28 proportion of social interaction time bin #2 vs #3)<br>p = 0.01 (MF P28 proportion of social interaction time bin #2 vs #4)*<br>p = 0.02 (MF P28 proportion of social interaction time bin #2 vs #5)*<br>p = 0.008 (MF P28 proportion of social interaction time bin #2 vs #6)**<br>p = 0.99 (MF P28 proportion of social interaction time bin #3 vs #4)<br>p = 1 (MF P28 proportion of social interaction time bin #3 vs #5) | $\bar{x}$ = 0.29, sd = 0.36 (adult MM time bin #2 proportion of USVs)<br>$\bar{x}$ = 0.13, sd = 0.2 (adult MM time bin #2 proportion of social interaction)<br>$\bar{x}$ = 0.09, sd = 0.07 (adult MM time bin #3 proportion of USVs)<br>$\bar{x}$ = 0.04, sd = 0.07 (adult MM time bin #3 proportion of social interaction)<br>$\bar{x}$ = 0.05, sd = 0.07 (adult MM time bin #4 proportion of USVs)<br>$\bar{x}$ = 0.07, sd = 0.1 (adult MM time bin #4 proportion of social interaction)<br>$\bar{x}$ = 0.04, sd = 0.05 (adult MM time bin #5 proportion of USVs)<br>$\bar{x}$ = 0.07, sd = 0.07 (adult MM time bin #5 proportion of social interaction)<br>$\bar{x}$ = 0.06, sd = 0.08 (adult MM time bin #6 proportion of USVs)<br>$\bar{x}$ = 0.1, sd = 0.12 (adult MM time bin #6 proportion of social interaction) |

| | | | |
|---|---|---|---|
| | | | p = 0.95 (MF P28 proportion of social interaction time bin #3 vs #6)<br>p = 1 (MF P28 proportion of social interaction time bin #4 vs #5)<br>p = 1 (MF P28 proportion of social interaction time bin #4 vs #6)<br>p = 1 (MF P28 proportion of social interaction time bin #5 vs #6)<br>p = 0.97 (MF adult proportion of social interaction main effect of time bin)<br>p = 0.05 (FF P24 proportion of social interaction main effect of time bin)*<br>p = 0.06 (FF P24 proportion of social interaction time bin #1 vs #2)<br>p = 0.13 (FF P24 proportion of social interaction time bin #1 vs #3)<br>p = 0.21 (FF P24 proportion of social interaction time bin #1 vs #4)<br>p = 0.21 (FF P24 proportion of social interaction time bin #1 vs #5)<br>p = 0.04 (FF P24 proportion of social interaction time bin #1 vs #6)*<br>p = 1 (FF P24 proportion of social interaction time bin #2 vs #3)<br>p = 0.99 (FF P24 proportion of social |

| | | | |
|---|---|---|---|
| | | | interaction time bin #2 vs #4)<br>p = 0.99 (FF P24 proportion of social interaction time bin #2 vs #5)<br>p = 1 (FF P24 proportion of social interaction time bin #2 vs #6)<br>p = 1 (FF P24 proportion of social interaction time bin #3 vs #4)<br>p = 1 (FF P24 proportion of social interaction time bin #3 vs #5)<br>p = 0.99 (FF P24 proportion of social interaction time bin #3 vs #6)<br>p = 1 (FF P24 proportion of social interaction time bin #4 vs #5)<br>p = 0.96 (FF P24 proportion of social interaction time bin #4 vs #6)<br>p = 0.96 (FF P24 proportion of social interaction time bin #5 vs #6)<br>p < 0.001 (FF P28 proportion of social interaction main effect of time bin)**<br>p < 0.001 (FF P28 proportion of social interaction time bin #1 vs #2)**<br>p < 0.001 (FF P28 proportion of social interaction time bin #1 vs #3)**<br>p < 0.001 (FF P28 proportion of social interaction time bin #1 vs #4)** |

| | | | |
|---|---|---|---|
| | | | $p < 0.001$ (FF P28 proportion of social interaction time bin #1 vs #5)** $p < 0.001$ (FF P28 proportion of social interaction time bin #1 vs #6)** $p = 0.87$ (FF P28 proportion of social interaction time bin #2 vs #3) $p = 1$ (FF P28 proportion of social interaction time bin #2 vs #4) $p = 1$ (FF P28 proportion of social interaction time bin #2 vs #5) $p = 0.34$ (FF P28 proportion of social interaction time bin #2 vs #6) $p = 0.79$ (FF P28 proportion of social interaction time bin #3 vs #4) $p = 0.94$ (FF P28 proportion of social interaction time bin #3 vs #5) $p = 0.93$ (FF P28 proportion of social interaction time bin #3 vs #6) $p = 1$ (FF P28 proportion of social interaction time bin #4 vs #5) $p = 0.26$ (FF P28 proportion of social interaction time bin #4 vs #6) $p = 0.44$ (FF P28 proportion of social interaction time bin #5 vs #6) $p < 0.001$ (FF adult proportion of social |

| | | | |
|---|---|---|---|
| | | | interaction main effect of time bin)** p < 0.001 (FF adult proportion of social interaction time bin #1 vs #2)** p < 0.001 (FF adult proportion of social interaction time bin #1 vs #3)** p < 0.001 (FF adult proportion of social interaction time bin #1 vs #4)** p < 0.001 (FF adult proportion of social interaction time bin #1 vs #5)** p < 0.001 (FF adult proportion of social interaction time bin #1 vs #6)** p = 0.88 (FF adult proportion of social interaction time bin #2 vs #3) p = 0.91 (FF adult proportion of social interaction time bin #2 vs #4) p = 1 (FF adult proportion of social interaction time bin #2 vs #5) p = 1 (FF adult proportion of social interaction time bin #2 vs #6) p = 1 (FF adult proportion of social interaction time bin #3 vs #4) p = 0.86 (FF adult proportion of social interaction time bin #3 vs #5) p = 0.87 (FF adult proportion of social interaction time bin #3 vs #6) |

| | | | |
|---|---|---|---|
| | | | p = 0.89 (FF adult proportion of social interaction time bin #4 vs #5)<br>p = 0.9 (FF adult proportion of social interaction time bin #4 vs #6)<br>p = 1 (FF adult proportion of social interaction time bin #5 vs #6)<br>p < 0.001 (MM P24 proportion of social interaction main effect of time bin)**<br>p = 0.01 (MM P24 proportion of social interaction time bin #1 vs #2)*<br>p < 0.001 (MM P24 proportion of social interaction time bin #1 vs #3)**<br>p < 0.001 (MM P24 proportion of social interaction time bin #1 vs #4)**<br>p < 0.001 (MM P24 proportion of social interaction time bin #1 vs #5)**<br>p < 0.001 (MM P24 proportion of social interaction time bin #1 vs #6)**<br>p = 0.92 (MM P24 proportion of social interaction time bin #2 vs #3)<br>p = 0.19 (MM P24 proportion of social interaction time bin #2 vs #4)<br>p = 0.6 (MM P24 proportion of social interaction time bin #2 vs #5)<br>p = 0.15 (MM P24 proportion of social |

| | | | |
|---|---|---|---|
| | | | interaction time bin #2 vs #6)<br>$p = 0.74$ (MM P24 proportion of social interaction time bin #3 vs #4)<br>$p = 0.99$ (MM P24 proportion of social interaction time bin #3 vs #5)<br>$p = 0.65$ (MM P24 proportion of social interaction time bin #3 vs #6)<br>$p = 0.98$ (MM P24 proportion of social interaction time bin #4 vs #5)<br>$p = 1$ (MM P24 proportion of social interaction time bin #4 vs #6)<br>$p = 0.95$ (MM P24 proportion of social interaction time bin #5 vs #6)<br>$p < 0.001$ (MM P28 proportion of social interaction main effect of time bin)**<br>$p < 0.001$ (MM P28 proportion of social interaction time bin #1 vs #2)**<br>$p < 0.001$ (MM P28 proportion of social interaction time bin #1 vs #3)**<br>$p < 0.001$ (MM P28 proportion of social interaction time bin #1 vs #4)**<br>$p < 0.001$ (MM P28 proportion of social interaction time bin #1 vs #5)**<br>$p < 0.001$ (MM P28 proportion of social interaction time bin #1 vs #6)** |

| | | | |
|---|---|---|---|
| | | | p = 0.99 (MM P28 proportion of social interaction time bin #2 vs #3)<br>p = 1 (MM P28 proportion of social interaction time bin #2 vs #4)<br>p = 1 (MM P28 proportion of social interaction time bin #2 vs #5)<br>p = 0.95 (MM P28 proportion of social interaction time bin #2 vs #6)<br>p = 0.99 (MM P28 proportion of social interaction time bin #3 vs #4)<br>p = 0.95 (MM P28 proportion of social interaction time bin #3 vs #5)<br>p = 1 (MM P28 proportion of social interaction time bin #3 vs #6)<br>p = 1 (MM P28 proportion of social interaction time bin #4 vs #5)<br>p = 0.96 (MM P28 proportion of social interaction time bin #4 vs #6)<br>p = 0.87 (MM P28 proportion of social interaction time bin #5 vs #6)<br>p < 0.001 (MM adult proportion of social interaction main effect of time bin)**<br>p < 0.001 (MM adult proportion of social interaction time bin #1 vs #2)**<br>p < 0.001 (MM adult proportion of |

| | | | |
|---|---|---|---|
| | | | social interaction time bin #1 vs #3)** p < 0.001 (MM adult proportion of social interaction time bin #1 vs #4)** p < 0.001 (MM adult proportion of social interaction time bin #1 vs #5)** p < 0.001 (MM adult proportion of social interaction time bin #1 vs #6)** p = 0.89 (MM adult proportion of social interaction time bin #2 vs #3) p = 0.98 (MM adult proportion of social interaction time bin #2 vs #4) p = 0.96 (MM adult proportion of social interaction time bin #2 vs #5) p = 1 (MM adult proportion of social interaction time bin #2 vs #6) p = 1 (MM adult proportion of social interaction time bin #3 vs #4) p = 1 (MM adult proportion of social interaction time bin #3 vs #5) p = 0.99 (MM adult proportion of social interaction time bin #3 vs #6) p = 1 (MM adult proportion of social interaction time bin #4 vs #5) p = 1 (MM adult proportion of social interaction time bin #4 vs #6) |

| | | | |
|---|---|---|---|
| | | | p = 1 (MM adult proportion of social interaction time bin #5 vs #6) |
